# Conformational sensors and domain-swapping reveal structural and functional differences between β-arrestin isoforms

**DOI:** 10.1101/725622

**Authors:** Eshan Ghosh, Hemlata Dwivedi, Mithu Baidya, Ashish Srivastava, Punita Kumari, Tomek Stepniewski, Hee Ryung Kim, Mi-Hye Lee, Jaana van Gastel, Madhu Chaturvedi, Debarati Roy, Shubhi Pandey, Jagannath Maharana, Ramon Guixà-Gonzàlez, Louis M. Luttrell, Ka Young Chung, Somnath Dutta, Jana Selent, Arun K. Shukla

## Abstract

Desensitization, signaling and trafficking of G protein-coupled receptors (GPCRs) are critically regulated by multifunctional adaptor proteins, β-arrestins (βarrs). The two isoforms of βarrs (βarr1 and 2) share a high degree of sequence and structural similarity, still however, they often mediate distinct functional outcomes in the context of GPCR signaling and regulation. A mechanistic basis for such a functional divergence of βarr isoforms is still lacking. Using a set of complementary approaches including antibody fragment based conformational sensors, we discover structural differences between βarr1 and 2 upon their interaction with activated and phosphorylated receptors. Interestingly, domain swapped chimeras of βarrs display robust complementation in functional assays thereby, linking the structural differences between the receptor-bound βarr1 and 2 with their divergent functional outcomes. Our findings reveal important insights into the ability of βarr isoforms to drive distinct functional outcomes, and underscore the importance of integrating this aspect in the current framework of biased agonism.

## INTRODUCTION

G Protein-Coupled Receptors (GPCRs), also referred to as seven transmembrane receptors (7TMRs), constitute a large family of integral membrane proteins in the human genome (Bjarnadottir et al., 2006) and a major class of drug targets (Santos et al., 2017). Upon activation by agonists, GPCRs couple to heterotrimeric G proteins followed by the generation of second messengers and downstream signaling. Subsequently, they are phosphorylated in their carboxyl-terminus and intracellular loops, which then results in coupling of β-arrestins (βarrs). Binding of βarrs interferes with further coupling of G proteins through steric hindrance, at least at the plasma membrane, and leads to receptor desensitization. Interestingly, βarrs also serve as adaptors for the components of clathrin machinery to mediate receptor endocytosis (Goodman et al., 1996; Laporte et al., 1999), and they can also scaffold various kinases to initiate several signaling pathways (DeWire et al., 2007; Luttrell et al., 1999; Luttrell et al., 2001; McDonald et al., 2000).

There are two isoforms of βarrs, referred to as βarr1 and 2 (also known as arrestin 2 and 3, respectively) sharing approximately 75% sequence identity, and both display an overall similar three-dimensional structure (Gurevich and Gurevich, 2015). For most GPCRs, both isoforms of βarrs are typically recruited to the receptor upon agonist stimulation, and participate in desensitization, endocytosis and signaling. Emerging data suggest that there exists a significant level of functional divergence between βarr1 and 2 for most GPCRs, and in some cases, they even display functional antagonism with each other (Figure 1A) (Ahn et al., 2003; Hara et al., 2011; Srivastava et al., 2015). Interestingly, the functional divergence of βarr isoforms is also manifested at the level of physiological outcomes downstream of several GPCRs (Srivastava et al., 2015; Trivedi et al., 2013; Walters et al., 2009)

**FIGURE 1.**
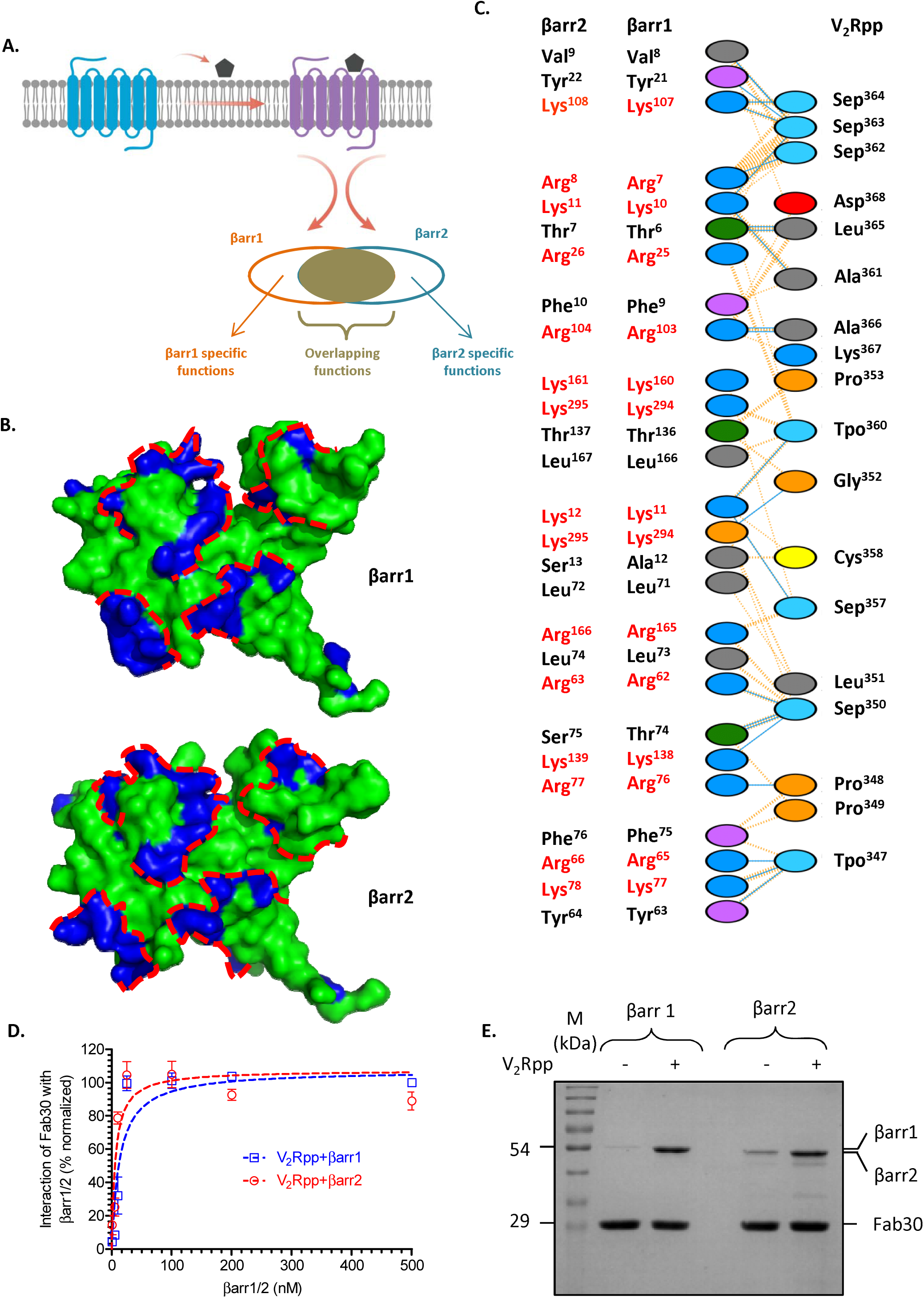
Overall structural and conformational similarity between V_2_Rpp-bound βarr1 and 2. **(A)** Most GPCRs typically recruit both isoforms of βarrs i.e. βarr1 and 2. The two isoforms typically mediate some overlapping functions but they also often display differential contributions in receptor desensitization, trafficking and signaling outcomes. **(B)** Lys/Arg residues on the N-domain of βarr1 are aligned into a spatial groove, and V_2_Rpp docks into this groove. Dotted red lines are manually drawn to highlight the groove where V_2_Rpp docks in the crystal structure. Corresponding Lys/Arg residues in βarr2 which are conserved at the primary sequence level, also exhibit a similar spatial arrangement as visualized using the crystal structure of βarr2 (PDB ID: 3P2D). **(C)** Schematic representation of amino acids in βarr1 that are within the interacting distance of V_2_Rpp based on previously determined crystal structure (PDB ID: 4JQI). The coordinates of the amino acid residues involved in binding with V_2_Rpp were submitted into PDBSum, and the interactions were mapped as a simplified ladder. Corresponding residues in βarr2 are listed to highlight the conserved nature of V_2_Rpp interacting residues in βarr1 and 2. Lys and Arg residues which form ionic interactions with the phosphate groups on V_2_Rpp are highlighted in red. Sep and Tpo represent phosphorylated serine and threonine residues in V_2_Rpp, respectively. Dotted blue and orange lines denote hydrogen bonding and non-bonded contacts, respectively. The residues have been colored according to their types, viz. blue - positive, red - negative, green - neutral, purple - aromatic, grey - aliphatic, orange - Pro and Gly and yellow - cysteine. **(D)** Fab30 recognizes V_2_Rpp-bound βarr1 and 2 to a comparable level. This experiment is carried out using an ELISA-based approach to measure the reactivity of Fab30 towards V_2_Rpp-bound βarr1 and 2 as described in the method section. Data represent average±SEM of three independent experiments, each carried out in duplicate. Data are normalized with respect to maximum signal obtained for V_2_Rpp-βarr1 condition (treated as 100%). **(E)** Similar reactivity of Fab30 towards V_2_Rpp-bound βarr1 and 2 is further confirmed by a co-IP experiment. A representative image of three independent experiments is shown here, and densitometry based quantification of all three experiments is presented in Figure S3A. See also Figure S1.

The functional divergence of βarr isoforms has direct implications for the conceptual framework of βarr-dependent signaling, biased-agonism, and in particular, for the development of βarr-biased ligands as novel GPCR therapeutics (Shukla et al., 2011). Still however, the mechanistic basis of their functional divergence is currently lacking, and it represents a key knowledge gap in our current understanding of GPCR-βarr interaction and signaling. Receptor-βarr interaction is a biphasic process involving the phosphorylated carboxyl-terminus (i.e. receptor tail) and the transmembrane bundle (i.e. receptor core) (Gurevich and Gurevich, 2004; Shukla et al., 2014). These two set of interactions result in the formation of partially-engaged (i.e. tail-engaged) and fully-engaged (i.e. core-engaged) receptor-βarr complexes, respectively. As recent studies have revealed distinct functional outcomes associated with these two conformations of receptor-βarr complexes (Cahill et al., 2017; Kumari et al., 2016; Kumari et al., 2017; Sente et al., 2018), we envisioned that key determinants of the functional divergence of βarr isoforms may lie at the level of structural and conformational differences between receptor-bound βarrs.

Accordingly, we set out to probe the conformations of receptor-bound βarrs using a battery of complementary approaches including biochemical and functional assays, synthetic antibody-based conformational biosensors, single particle electron microscopy, bimane fluorescence spectroscopy and molecular dynamics simulation. We discover potential structural differences between receptor-bound βarr1 and 2 and identify key regions in βarrs that are critical for imparting these differences. Using a domain-swapped chimera of βarrs, we also observe that the structural differences between βarr1 and 2 manifests in their distinct functional contributions downstream of GPCRs. Our findings provide important insights into GPCR-βarr interaction and they have direct implications for refining the framework of biased agonism at GPCRs.

## RESULTS

### Sequence and structural analysis of βarrs

The interaction of phosphorylated receptor tail (i.e. the carboxyl-terminus) with βarrs results in the formation of a partially-engaged complex. Crystal structure of βarr1 in complex with a phosphopeptide corresponding to the carboxyl-terminus of the human vasopressin receptor (V_2_R) (referred to as V_2_Rpp) has revealed the interaction interface between receptor-attached phosphate groups and positively charged residues in the N-domain of βarr1 (Shukla et al., 2013). Here, V_2_Rpp serves as a surrogate for the phosphorylated receptor tail and therefore, V_2_Rpp-βarr1 complex represents a close proxy of the partially-engaged receptor-βarr1 complex. Our analysis of the phosphate-interacting residues on βarr1 in this crystal structure, and their spatial surface mapping, revealed a groove along the N-domain of βarr1 that constitutes the docking interface for V_2_Rpp through a number of charge-charge interactions (Figure 1B, upper panel). As V_2_Rpp binds both βarr1 and 2 with comparable affinities (Nobles et al., 2007; Xiao et al., 2004), we analyzed the sequence and the three-dimensional structure of βarr2 to assess whether the spatial arrangement of phosphate-interacting residues may be conserved in both isoforms. Indeed, we not only observed that phosphate-interacting residues are highly conserved in βarr2 but their spatial distribution on the N-domain of βarr2 also forms a groove identical to βarr1 (Figure 1B, lower panel, Figure 1C and Figure S1). This observation suggests potentially similar binding mechanism during the first step of receptor-βarr interaction, and therefore, provides a rationale to probe it experimentally.

### Fab30 as a sensor of βarr activation reveals overall similarity between V_2_Rpp-bound βarr1 and 2

Based on sequence and structural analysis, we conceived that the overall conformation of partially-engaged βarr1 and 2 (i.e. in complex with V_2_Rpp) may be similar. To probe this, we used a synthetic antibody fragment (referred to as Fab30) as a sensor of βarr conformation. Fab30 selectively recognizes active βarr1 in both, the partially-engaged and the fully-engaged complexes with the receptor (Kumari et al., 2016; Kumari et al., 2017; Shukla et al., 2013; Shukla et al., 2014). As the ability of Fab30 to recognize βarr2 has not been evaluated previously, we first identified the paratope residues for Fab30 binding on βarr1 based on the crystal structure of V_2_Rpp-βarr1-Fab30 complex (Shukla et al., 2013), and confirmed that they are mostly conserved in βarr2 (Figure S2A-C). Thus, Fab30 should recognize V_2_Rpp-bound βarr2 as well and in fact, we observed a robust interaction of Fab30 with V_2_Rpp-bound βarr2 at comparable levels to βarr1 in two parallel assays based on ELISA and co-immunoprecipitation (co-IP) (Figure 1D-E and Figure S3A). In addition, a single chain variable fragment version of Fab30, referred to as ScFv30, also exhibited a similar pattern of reactivity towards V_2_Rpp-boundβarr1 and 2 (Figure S3B-C). Under similar experimental conditions, a control Fab (Fab-CTL) that does not recognize βarr1 failed to exhibit any significant specific binding in co-IP experiments (Figure S3D-E). These data suggest that the overall conformation of βarr1 and 2 in complex with V_2_Rpp, as detected by Fab30 reactivity, are similar.

### Fab30 reactivity suggests potential conformational differences between receptor-bound βarr1 and 2

We next set out to measure the reactivity of Fab30 towards activated and phosphorylated receptor-bound βarr1 and 2. Here, we used two different GPCRs, the V_2_R and a chimeric β2-adrenergic receptor, referred to as β_2_V_2_R, where the carboxyl-terminus of the β_2_AR is replaced with that of the V_2_R (Oakley et al., 2000; Thomsen et al., 2016). Although we have previously reported that Fab30 robustly recognizes receptor-βarr1 complexes (Kumari et al., 2016; Shukla et al., 2014), in order to further establish Fab30 as a reliable sensor of receptor-bound βarr conformation, we measured its reactivity towards β_2_V_2_R-βarr1 complexes formed in response to a set of ligands having different efficacies ranging from inverse agonists, partial agonists to full agonists. We observed an excellent correlation between Fab30 reactivity (measured using co-IP assay) and the relative efficacy of ligands as measured by their cAMP response (Figure S4A-E). This observation underlines the ability of Fab30 to report a pharmacologically relevant receptor-bound βarr1 conformation and therefore, allows us to compare the conformation of receptor-bound βarr1 and 2.

We employed two parallel approaches based on ELISA and co-IP assays using activated and phosphorylated β_2_V_2_R and V_2_R that were generated following a previously published protocol (Kumari et al., 2016; Kumari et al., 2017). As expected, Fab30 robustly recognized receptor-bound βarr1, surprisingly however, it failed to recognize receptor-bound βarr2 for both, the β_2_V_2_R (Figure 2A-B, Figure S5A) and the V_2_R (Figure S5B-C). ScFv30 also exhibits a pattern identical to Fab30 i.e. it recognizes receptor-bound βarr1 but not βarr2 (Figure S5D-E). As the key residues in βarr1 responsible for binding Fab30 are mostly conserved in βarr2, and Fab30 can robustly bind V_2_Rpp-βarr2 complex, the lack of Fab30 reactivity towards receptor-bound βarr2 is unlikely to result from differences in its interaction interface between βarr1 and 2. In agreement with previous studies (Oakley et al., 2000), we also observed that β_2_V_2_R and V_2_R robustly interact with βarr2 (Figure S6A-C), and therefore, the lack of Fab30 reactivity is also not because of the inability of β_2_V_2_R to bind βarr2.

**FIGURE 2.**
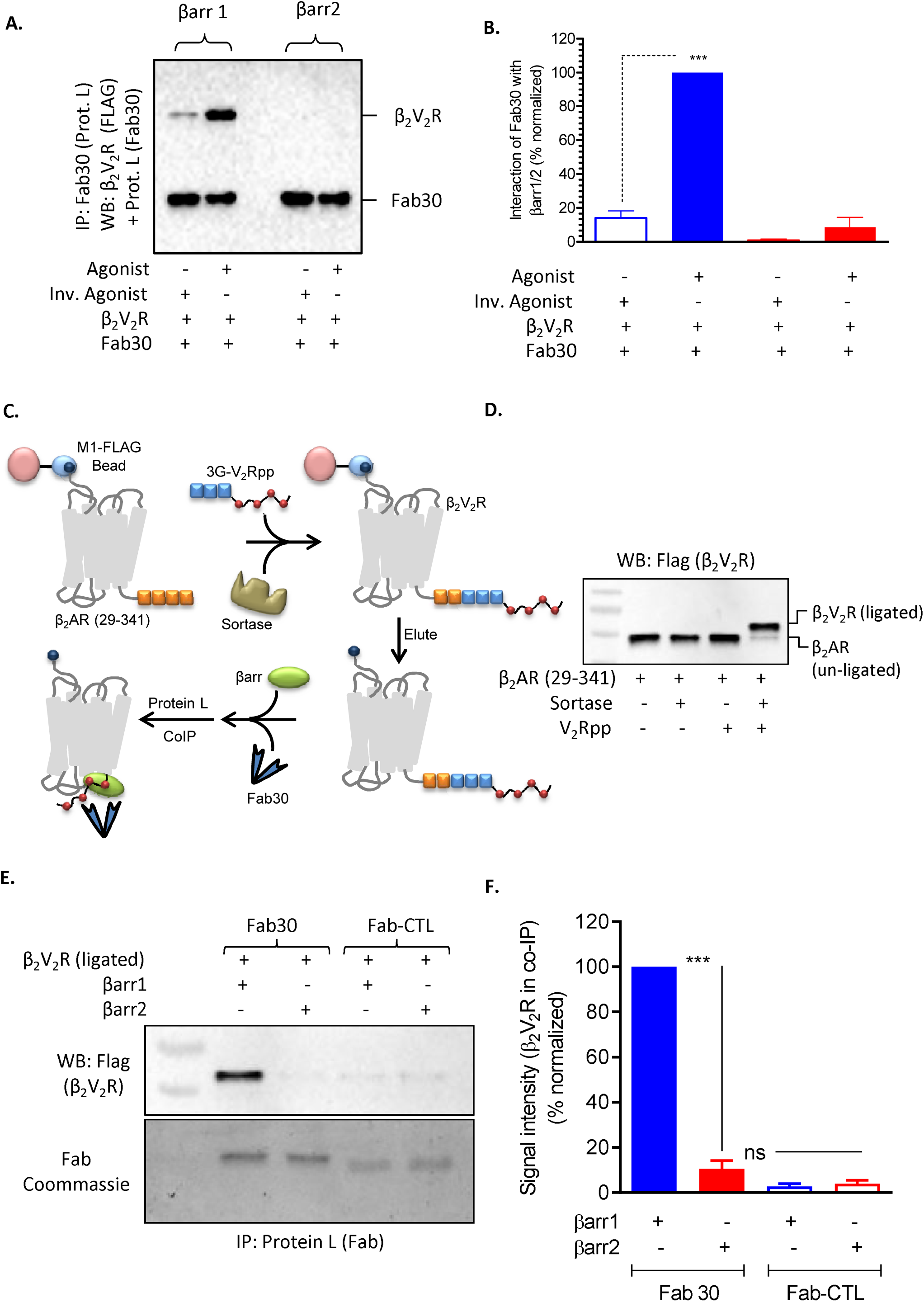
Fab30 reactivity pattern reveals potential conformational differences between receptor-bound βarr1 and 2. **(A)** Fab30 robustly recognizes β_2_V_2_R-bound βarr1 but not βarr2 as assessed by a co-IP assay followed by visualization using Western blot. Here, Carazolol (1μM) is used as an inverse agonist and BI-167107 (1μM) is used as an agonist to stimulate cells expressing FLAG-tagged β_2_V_2_R. A representative image from three independent experiments is shown. **(B)** Densitometry-based quantification of data presented in panel A. Values represent mean±SEM of three independent experiments analyzed using One-Way ANOVA with Bonferroni post-test (***P<0.001). Data are normalized with respect to agonist-β_2_V_2_R-βarr1 condition (treated as 100%). **(C)** Schematic flow-chart of sortase-based chemical ligation of V_2_Rpp with truncated β_2_AR (29-341), and subsequent co-IP experiment to measure the reactivity of Fab30 towards homogenously phosphorylated receptor. **(D)** Efficiency of sortase-based ligation of V_2_Rpp to β_2_AR (29-341) as measured by Western blotting. A representative blot from two independent experiments is shown. **(E)** Fab30 fails to recognize βarr2 in complex with homogenously phosphorylated β_2_V_2_R. After sortase-based chemical ligation of V_2_Rpp, the resulting β_2_V_2_R was incubated with equal concentrations of βarr1/2, and Fab30/Fab-CTL followed by co-immunoprecipitation using Protein L beads. The reactivity of Fab30 with receptor-bound βarr1/2 was evaluated by Western blot. **(F)** Densitometry-based quantification of Fab30 reactivity towards receptor-bound βarr1 and 2 as measured in panel E. Data represent average±SEM of three independent experiments, normalized with respect to βarr1 (treated as 100%). See also Figures S2-7.

In order to rule out the affinity difference of Fab30 for βarr1 vs. βarr2, we carried out titration co-IP experiments, first with increasing concentrations of V_2_Rpp, and second, with increasing concentration of β_2_V_2_R while keeping the concentrations of βarr1 and βarr2 constant. Fab30 recognizes V_2_Rpp-bound βarr2 as efficiently as V_2_Rpp-bound βarr1, even at partial occupancy of V_2_Rpp (Figure S7A-B). Moreover, even at nine-fold higher concentration of β_2_V_2_R, we still did not observe any detectable reactivity of Fab30 towards receptor-bound βarr2 (Figure S7C). These data suggest that the lack of Fab30 reactivity towards receptor-bound βarr2 is not due to an affinity difference of Fab30 or available stoichiometry of phosphorylated tail between V_2_Rpp vs. β_2_V_2_R experiments.

In order to validate our data in cellular context, we next expressed HA-tagged intrabody version of Fab30 (referred to as Ib30) together with β_2_V_2_R and either βarr1 or βarr2 in HEK-293 cells followed by a co-IP experiment. Similar to *in-vitro* experiments performed with purified proteins, we found that even in the cellular context, intrabody 30 recognizes receptor-bound βarr1 but not βarr2 (Figure S7D). Taken together, these findings suggest that there are potential conformational differences between βarr1 and 2 in complex with activated and phosphorylated receptors, which in turn results in the lack of Fab30 reactivity towards receptor-βarr2 complex.

### Fab30 reactivity towards βarr1 and 2 in presence of homogenously phosphorylated receptor

For the experiments mentioned in Figure 2A-B and Figure S5, we have utilized *in-cellulo* phosphorylated receptors. A potential concern may be heterogeneous phosphorylation of the receptor carboxyl-terminus when compared to synthetic V_2_Rpp with well-defined phosphorylation pattern leading to the lack of Fab30 reactivity towards receptor-bound βarr2. To rule out this possibility, we employed two parallel approaches. First, we used Sortase enzyme-based chemical ligation of V_2_Rpp to truncated β_2_AR (29-341) in order to generate a chimeric β_2_V_2_R with well-defined and homogenous phosphorylation pattern, identical to that present in V_2_Rpp (Figure 2C-D). Interestingly, we observed that similar to *in-cellulo* phosphorylated receptor, this chemically ligated version of the receptor also induces a conformation in βarr2 that is not recognized by Fab30 (Figure 2E-F). Second, we generated a series of V_2_R phosphorylation mutants lacking either the individual phosphorylation sites or cluster of phosphorylation sites, and compared the ability of Fab30 to recognize receptor-bound βarr1 and 2 for these mutants. Our notion was that if the lack of Fab30 reactivity towards receptor-bound βarr2 arises from the contribution of phosphorylation at a specific site (or lack thereof), we should be able to gain the reactivity in one (or several) of these mutants. However, we did not observe the gain of Fab30 reactivity in any of these receptor mutants (Figure S8A-C). Taken together, these data suggest that the lack of Fab30 recognition for receptor-bound βarr2 does not arise from heterogeneous or site-specific phosphorylation of the receptor.

### Corroborating evidence for potential conformations differences between βarr1 and 2

Leading up to this point, our data based on Fab30 recognition suggest potential conformational difference between receptor-bound βarr1 and 2. In order to corroborate these findings further, we tested a series of additional Fabs that we have recently generated and characterized to selectively recognize V_2_Rpp-bound βarr1 (Ghosh et al., 2017). Similar to Fab30, these additional Fabs also interacted comparably with V_2_Rpp-bound βarr1 and 2 (Figure S8D), and efficiently recognized the complex of βarr1 with activated and phosphorylated receptor (Figure S8E). Interestingly however, these additional Fabs also displayed no detectable recognition towards receptor-bound βarr2 (Figure S8E). Although the binding epitope of these additional Fabs on βarr1 has not been precisely determined yet, their reactivity pattern supports the notion of conformational differences between receptor-bound βarr isoforms.

As an additional line of evidence for conformational differences between receptor-bound βarr1 and 2, independent of Fab30 reactivity as readout, we employed bimane fluorescence spectroscopy. Here, we used purified βarr1 and 2 that are bimane labeled in their C-loop (βarr1^245C^ and βarr2^246C^) which forms a key interface for receptor interaction and exhibits conformational rearrangement during receptor interaction (Latorraca et al., 2018) (Figure 3A). Bimane is an environmentally sensitive fluorophore and its fluorescence intensity changes when it experiences a change in its environment or if it experiences a change in its proximity to a quenching residue. We observed a decrease in bimane fluorescence for βarr1 upon its interaction with the receptor while a significant increase for βarr2 (Figure 3B). These directionally opposite changes in bimane fluorescence intensities for βarr1 and 2 upon their interaction with the receptor suggest that their C-loops are positioned in different environments, and provide additional corroborating evidence for their conformational difference in receptor-bound states. We did not observe a significant change in bimane fluorescence intensity upon V_2_Rpp binding which can be interpreted to reflect conformational similarity between V_2_Rpp-bound βarr1 and 2 with respect to C-loop.

**FIGURE 3.**
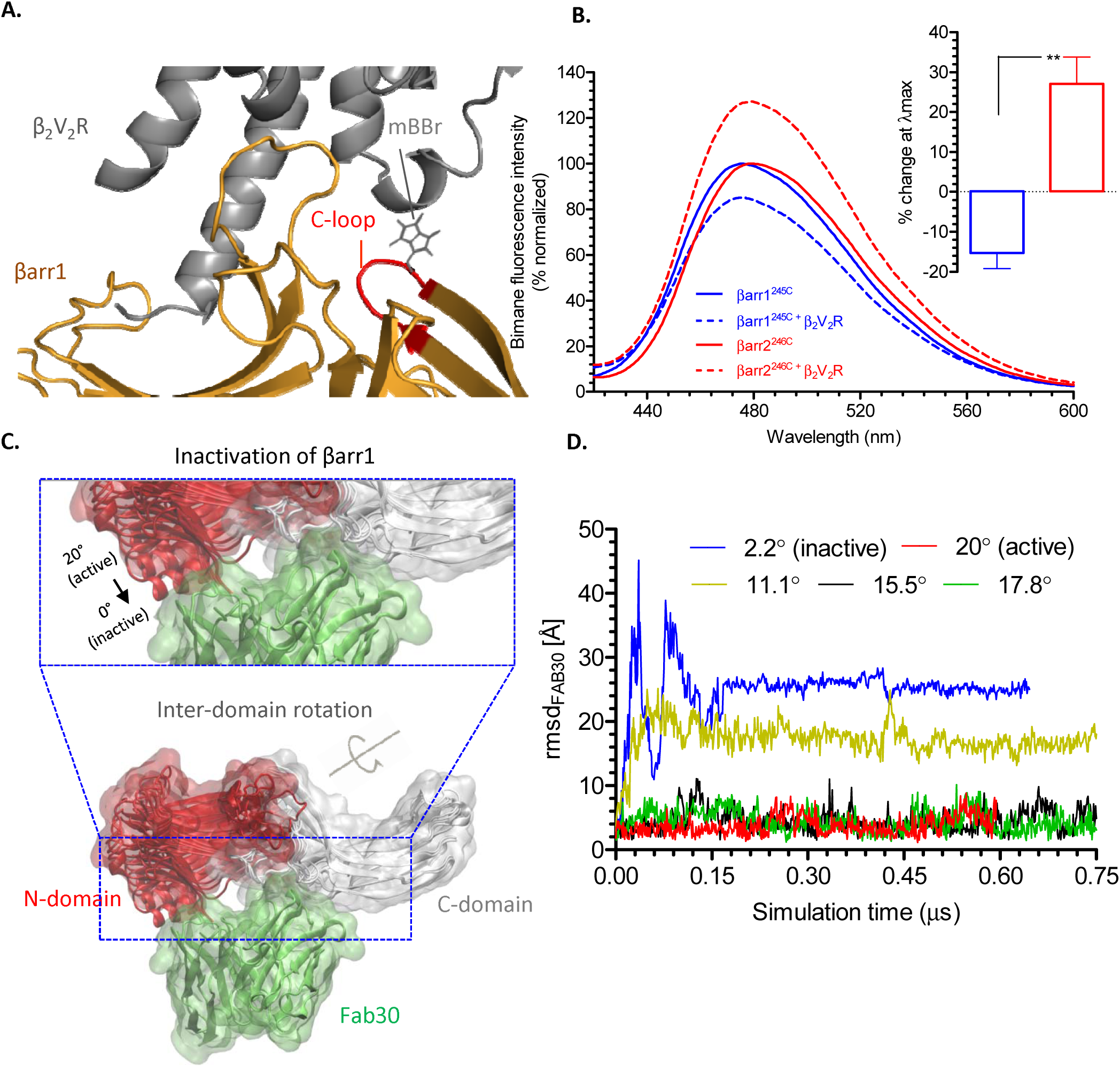
Fluorescence spectroscopy and molecular dynamics simulation provide insights into conformational differences between βarr isoforms. **(A)** Schematic representation of bimane fluorescence spectroscopy experiment where monobromobimane (mBBr) is chemically attached to a cysteine, engineered in the C-loop loop of βarrs. The fluorescence of mBBr is sensitive to its environment and therefore, a change in fluorescence intensity upon interaction of βarrs with the receptor reflects a conformational change. **(B)** The fluorescence intensity of βarr1^mBBr^ decreases upon its interaction with β V R while that of βarr2^mBBr^ increases significantly. Here, purified βarrs^mBBr^ were incubated with purified β_2_V_2_R (agonist-bound and phosphorylated) at a molar ratio of βarr1:β_2_V_2_R (1:3 in a concentration range of 1-5μM). As a reference, the fluorescence intensity of βarrs^mBBr^ alone was measured first and used for normalization (treated as 100%). The inset shows the differences in bimane fluorescence at λ_max_ for receptor-bound βarr1 and 2. **(C-D)** Fab30 binding stability depends on the inter-domain rotation angle of βarr1. The binding stability of the ScFv version of Fab30 (green surface) in complex with βarr1 (N-domain: red surface, C-domain: white surface) is measured as RMSD of Fab30 backbone atoms (RMSD_Fab30_) for different activation states of βarr1 (inter-domain rotation angles 2.2°, 11.1°, 15.5°, 17.8° and 20°). Inter-domain rotation angles 15.5° and 20° in βarr1 result in a stable RMSD_Fab30_ progression. Rotation angles of 2.2° and 11.1° provoke a rapid increase of the RMSD_Fab30_ (i.e. binding instability) during the first 50 ns of simulation time due to clashes of the N-domain with Fab30. See also Figures S9A.

### Structural insights into conformational differences between receptor-bound βarrs

In order to better understand Fab30 reactivity pattern and conformational differences between receptor-bound βarr1 and 2, we employed molecular dynamics simulation to gain structural and mechanistic insight. Crystal structure of βarr1 in complex with V_2_Rpp and Fab30 has revealed a major rotation of the C-domain relative to the N-domain by approximately 20° (Shukla et al., 2013). We postulated that this inter-domain rotation in βarrs may indeed be the primary determinant for effective recognition by Fab30, and in order to test this, we performed MD simulations monitoring the stability of Fab30 binding to βarr1 conformers with different inter-domain rotation angles in solution (Figure 3C-D and Figure S9A). Here, Fab30 binding stability is assessed as root-mean-square deviation (RMSD) of the backbone atoms along the simulation time. We found that Fab30 remains stable when bound to the conformers with a rotation angle >15° (i.e. stable RMSD progression) (Figure 3C-D). In contrast, a significant instability of Fab30 interaction is observed when it is bound to arrestin conformers with a rotation angle <15° (i.e. drastic increase in RMSD) (Figure 3C-D). Such instability is not surprising because the N-domain of βarr1 approaches Fab30 when the inter-domain rotation relaxes towards the inactive (or basal) state (i.e. rotation angle decreasing from 20° to 0°) which in turn results in unfavorable contacts and steric clashes (Figure 3C-D). In other words, Fab30 reactivity can be considered as readout of the degree of inter-domain rotation in βarrs upon activation, and therefore, it is plausible that the inter-domain rotation in receptor-bound βarr2 is significantly smaller than βarr1, resulting in lack of recognition by Fab30.

MD simulation data presented above raise the possibility that the occurrence of receptor core-engagement with βarr2 may be a critical determinant in driving its distinct receptor-bound conformation compared to βarr1. In fact, a comparison of the interaction of βarrs in the absence of “tail-engagement” using carboxyl-terminus truncated V_2_R (referred to as V_2_R^ΔC-term^) provides potential evidence for stronger “core interaction” for βarr2 compared to βarr1 (Figure S9B). This finding may be interpreted to suggest that receptor-βarr2 complex may have a predominant distribution in the “fully-engaged” conformation, which is different from that of βarr1, therefore, providing a plausible explanation for near-complete lack of Fab30 reactivity. However, in the absence of high-resolution structures of fully-engaged receptor-βarr complexes, it is not feasible to precisely determine the contribution of core-engagement towards imparting distinct βarr conformations, and future structural studies may illuminate the mechanism of distinct βarr conformations upon their interaction with the receptors.

### Distal C-domain in βarrs is important for the structural differences between the two isoforms

In order to identify the key regions in βarrs that are potentially responsible for imparting distinct conformations on βarr1 and 2, we generated a series of chimeric βarr constructs, and tested the ability of Fab30 to recognize their complexes with the receptor. Unlike visual-arrestins and βarr1, βarr2 lacks the c-edge loops (these are different from the C-loop described earlier) which are demonstrated to anchor the C-domain of visual-arrestin to the plasma membrane upon its recruitment to rhodopsin (Lally et al., 2017). Therefore, we first generated a βarr2 construct where we grafted the c-edge loop1 of βarr1 in the corresponding position of βarr2 and tested its reactivity to Fab30 in receptor-bound conformation (Figure S9C-E). However, similar to βarr2, this loop-grafted construct also failed to exhibit detectable recognition by Fab30 (Figure S9C-E) suggesting that the lack of c-edge loop1 in βarr2, and thereby, potential lack of membrane anchoring, is not responsible for its conformational difference with βarr1.

Next, we generated a construct (referred to as swap 1) harboring the N-domain of βarr1 and the C-domain of βarr2, and tested the ability of Fab30 to recognize it in receptor-bound conformation. Fab30 did not recognize receptor-bound swap 1 suggesting that it adopts a conformation similar to βarr2, and that the primary determinants of distinct conformations of receptor-bound βarr1 and 2 are likely encoded in the C-domain (Figure 4A-B and Figure S9F). This is further confirmed by a reverse chimera (referred to as swap 2) harboring the N-domain of βarr2 and the C-domain of βarr1 which is effectively recognized by Fab30 (Figure 4C-D and Figure S9G). A series of additional chimeric constructs (referred to as swap 3-5) harboring the N-domain of βarr2 and different segments of the C-domain of βarr1 reveal that the structural determinants of distinct conformations of receptor-bound βarr 1 and 2 primarily reside in the distal C-domain at the primary sequence level (Figure 4C-D). The pattern of Fab30 reactivity also indicates that the conformation of receptor-bound swap1 is similar to βarr2 while that of swap2 is similar to receptor-bound βarr1. To provide additional support for the interaction of swap2 with the receptor in a manner similar to βarr1, we carried out negative-staining based single particle EM analysis of Fab30 stabilized β_2_V_2_R-βarr^swap2^ complex (Figure 4E). The β_2_V_2_R-βarr^swap2^-Fab30 complex also exhibits a biphasic interaction with conformational distribution between the partially-engaged and fully-engaged complexes, similar to what is previously observed for the β_2_V_2_R-βarr1-Fab30 complex (Shukla et al., 2014). We also tested the ability of Ib30 to recognize receptor-bound swap2 in cellular context, and in agreement with the data presented in Figure 4C-D, Ib30 robustly recognizes receptor-bound swap2, at a level similar to that of βarr1 (Figure 5A-B). This observation further suggests an overall similar conformation adopted by receptor-bound βarr1 and swap2, and provides supporting evidence for the cellular relevance of the data obtained with chimeric βarr constructs *in-vitro*.

**FIGURE 4.**
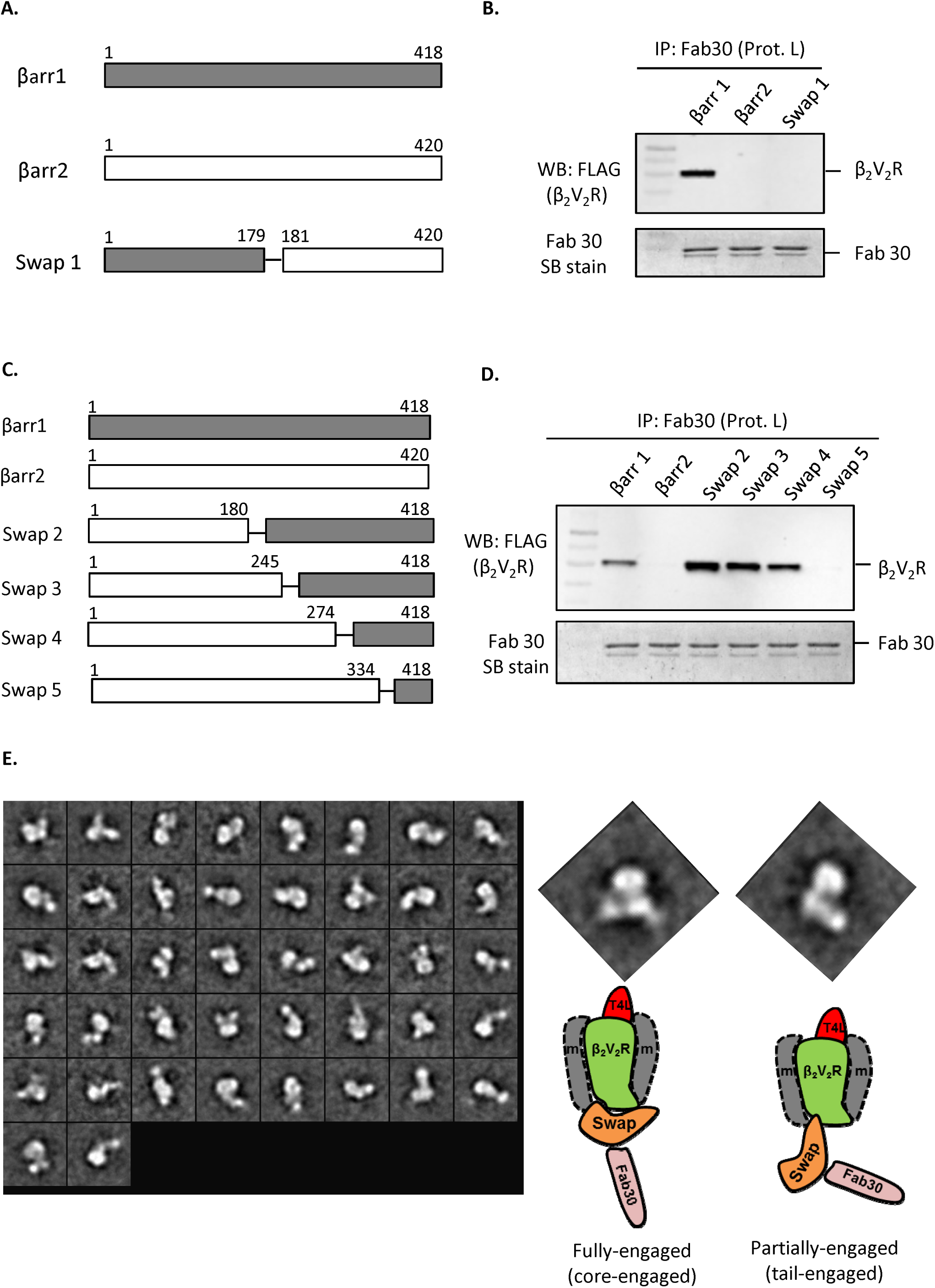
Identification of structural regions that impart conformational differences between receptor-bound βarr1 and 2. **(A)** Schematic representation of the swap1 construct that harbors the N-domain of βarr1 and the C-domain of βarr2. **(B)** Co-IP experiment reveals that Fab30 fails to detect receptor-bound conformation of swap1 indicating its conformational similarity with βarr2. Fab30 was mixed and incubated with purified (agonist-bound and phosphorylated) β_2_V_2_R and swap1 followed by co-IP. Samples were visualized by Western blotting (FLAG-β_2_V_2_R) or SimplyBlue staining (Fab30). A representative image of three independent experiments is shown here. **(C)** Schematic representation of swap2-5 constructs that harbor N-domain of βarr2 and different stretches of the C-domain of βarr1. (D) A co-IP experiment, carried out as described for panel B, reveals that the distal C-domain imparts conformational differences between βarr1 and 2. **(E)** Single particle negative staining EM analysis of the agonist-β_2_V_2_R-swap2-Fab30 complex. The agonist-β_2_V_2_R-swap2-Fab30 complex was reconstituted by mixing purified (agonist-bound and phosphorylated) β_2_V_2_R, swap2 and Fab30 proteins followed by size-exclusion chromatography. Subsequently, the complex preparation was immobilized on EM grids followed by negative-staining and visualization as described in the methods section. The right panel shows two representative 2-D class averages depicting the partially-engaged and fully-engaged complexes. Schematic representation of these two conformations is presented for the ease of visualization. See also Figure S9-10.

**FIGURE 5.**
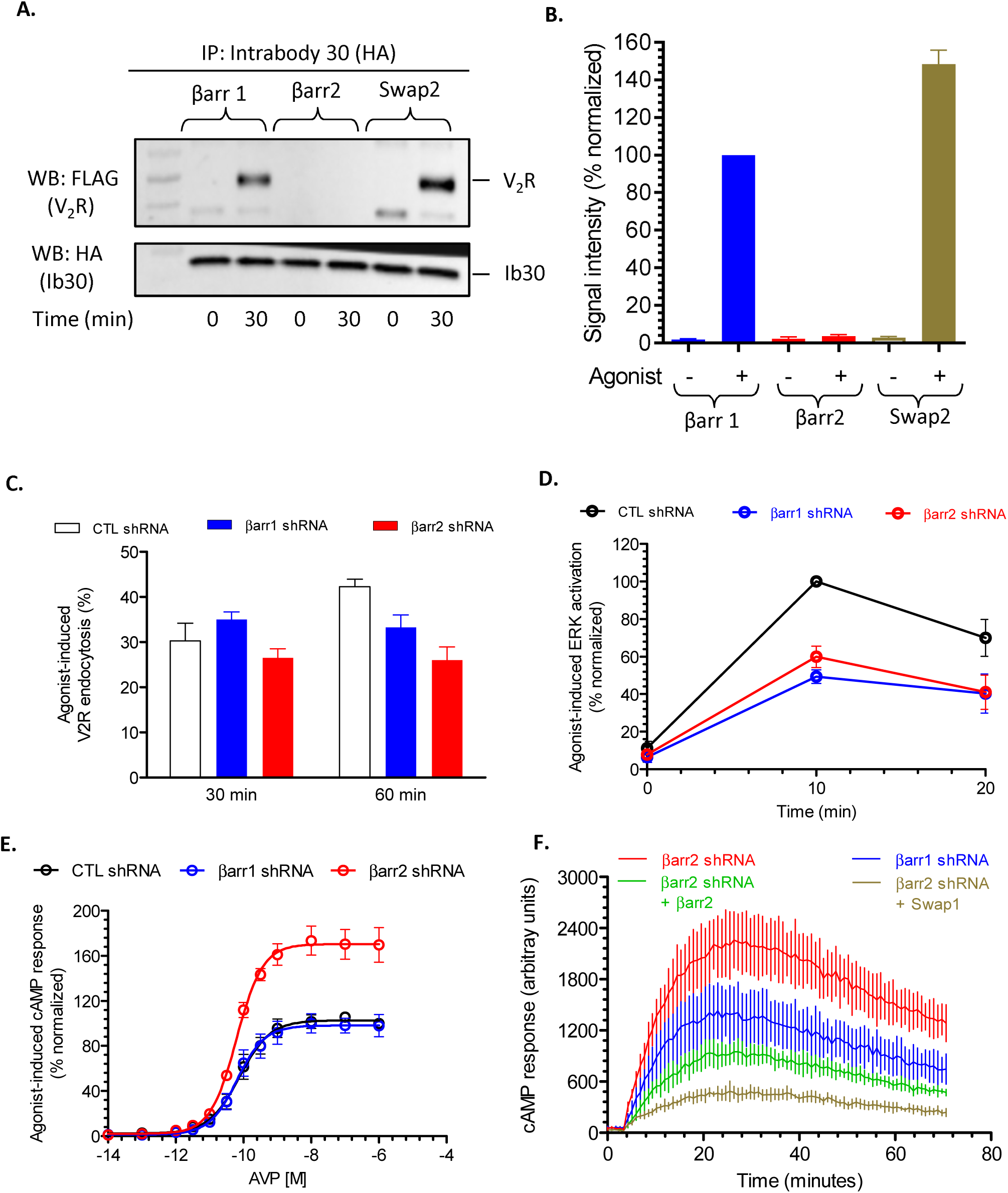
A domain-swapped chimera of βarrs gains Fab30 reactivity in cellular context and exhibits functional complementation. **(A)** HEK-293 cells expressing FLAG-V_2_R, βarr1, 2 or swap 2 and Ib30 were stimulated with AVP (100nM) for indicated time-points followed by co-IP using the HA tag on Ib30. The interaction of Ib30 with receptor-βarr complexes was visualized by Western blotting. A representative image from three independent experiments is shown. **(B)** Densitometry-based quantification of data presented in panel A. Values represent average signal intensity ± SEM normalized with respect to agonist-V_2_R-βarr1 condition (treated as 100%). **(C)** Agonist-induced endocytosis of V_2_R measured by whole cell ELISA in HEK-293 cells under control (CTL), βarr1 or βarr2 knock-down conditions. Data represent average±SEM of five independent experiments each carried out in duplicate. % endocytosis as measured by the surface level of V_2_R before and after agonist-stimulation is presented in the graph. **(D)** Agonist-induced ERK MAP kinase activation downstream of V_2_R as measured by Western blotting in cells under control (CTL), βarr1 or βarr2 knock-down conditions. Data represent average±SEM of five independent experiments and normalized with maximal ERK activation in CTL condition (treated as 100%). A representative image of these experiments is shown in Figure S10C. **(E)** Agonist-induced cAMP response in HEK-293 cells expressing V_2_R under control (CTL), βarr1 or βarr2 knock-down conditions using the Glo-sensor assay. Data represent average±SEM of three independent experiments each carried out in duplicate and normalized with the maximal cAMP response in CTL condition (treated as 100%). **(F)** Exogenous expression of βarr2 or swap1 lowers the enhanced levels of cAMP in βarr2 knock-down cells suggesting a potential link between receptor-bound βarr2 conformation and receptor desensitization. Here, the cells were stimulated with saturating concentration of agonist (100nM AVP) for indicated time-points. A representative profile from three independent experiments, each performed in duplicate, is shown here. See also Figure S10.

### A potential link between βarr conformations and their distinct functional contributions

To probe whether conformational differences between receptor-bound βarr isoforms may be directly linked to their functional divergence, we first measured the contribution of βarr1 and 2 in agonist-induced endocytosis, ERK MAP kinase activation and cAMP response for the V_2_R under βarr1 or 2 knockdown conditions (Figure 5C-F and Figure S10A). We found that the presence of either βarr1 or 2 is capable of supporting agonist-induced endocytosis of the V_2_R (Figure 5C) suggesting that the two isoforms are functionally redundant in mediating receptor endocytosis. Interestingly, agonist-induced ERK1/2 MAP kinase phosphorylation downstream of V_2_R is sensitive to the depletion of either isoforms of βarrs (Figure 5D and Figure S10C) suggesting that both isoforms are involved. Strikingly however, agonist-induced cAMP response is significantly enhanced upon βarr2 depletion (Figure 5E) indicating a predominant role of βarr2 in receptor desensitization compared to βarr1. The surface expression of V_2_R in βarr1 or 2 depleted cells are comparable to each other (Figure S10B). As expected, exogenous expression of βarr2 in βarr2 knock-down cells lowers the level of cAMP (Figure 5F). Most interestingly, exogenous expression of swap1, which is conformationally similar to βarr2, also effectively lowers the enhanced level of cAMP in βarr2 knock-down cells (Figure 5F). Taken together, these data suggest that receptor-bound βarr2 conformation is more effective in driving receptor desensitization compared to βarr1, and thus potentially connects the conformational differences between receptor-bound βarrs with their functional divergence.

## DISCUSSION

Most GPCRs recruit both βarr1 and 2 upon agonist stimulation, which in turn mediate and regulate receptor desensitization, endocytosis and signaling. Although both isoforms are individually capable of mediating the above-mentioned functions, interestingly, they often display differential contribution towards these functions and subsequent physiological outcomes for different receptor systems. This paradigm, now observed across multiple GPCRs (Srivastava et al., 2015), suggests potential differences at structural and conformational levels in receptor-bound states of βarr1 and 2. We observed that while the docking interface for the phosphorylated receptor tail and the resulting conformations in βarr1 and 2 are similar to each other, they appear to adopt different conformations upon their engagement with activated and phosphorylated receptors.

These findings suggest that a differential conformational rearrangement may happen in βarr1 vs. 2 when they transition from the partially-engaged to fully-engaged complex involving the receptor-core. In other words, the core-engagement between the receptor and βarrs may impart distinct structural changes in the two isoforms of βarrs. In fact, comparison of the crystal structures of pre-activated visual-arrestin (i.e. splice variant p44) with rhodopsin-visual-arrestin complex reveals significant structural changes in visual arrestin (Kang et al., 2015; Kim et al., 2013; Zhou et al., 2017). Moreover, a recent study on rhodopsin-visual-arrestin system has also suggested that both, the receptor-tail and the receptor-core are capable of inducing activating conformational changes in visual-arrestin, independent of each other (Latorraca et al., 2018). Additional studies in cellular context have also reported that the core-engagement can drive an active conformation in βarr2 which allows it to enrich in clathrin coated structures, even in the absence of a stable complex with the receptor (Eichel et al., 2018; Eichel et al., 2016). While these previous studies align with our findings raising the possibility of core-engagement driving the conformational differences between receptor-bound βarr1 and 2, future structural studies should illuminate the structural mechanism underlying this interesting phenomenon. As different GPCRs have diverse signatures of phosphorylatable residues in their carboxyl-terminus or in intracellular loops, and may have different levels of core-engagement, it may not be surprising to discover additional levels of conformational diversity in βarrs which makes this system precisely tunable in a context dependent manner (Ranjan et al., 2017).

It is also intriguing that the differences between βarr1 and 2 appear to arise primarily from the distal C-domain, a region that is not only most diverse between the two isoforms at the primary sequence level but also harbors the interface for several interaction partners such as clathrin, adaptin and TRAF6. Therefore, it is tempting to speculate that receptor-bound βarr1 and 2 may also differ in their ability to scaffold different partners, owing to their conformational differences. In fact, such a scenario is supported by a global interactomics analysis where a significant difference between the interactome of βarr1 and 2 is reported (Xiao et al., 2007). Still however, future studies are required to probe this interesting possibility further including a precise mapping of βarr residues that may be responsible for the difference of Fab30 reactivity between the βarr isoforms.

A previously published hydrogen/deuterium exchange (HDX) study of βarrs using pre-activated βarr mutants (βarr1^R169E^ and βarr2^R170E^) provides additional corroborating evidence for our findings (Yun et al., 2015). A re-analysis of this previous study reveals a significant difference between the HDX pattern of lariat loop peptides when we compare βarr1^WT^-βarr1^R169E^ and βarr2 ^WT^-βarr2^R170E^ with each other. For example, we observed a significantly higher rate of deuterium uptake in the 279-289 segment of βarr1 between the WT and R^169^E mutant but the corresponding region in βarr2 does not exhibit a significant difference between the wild-type and R^170^E mutant (Figure S11). Although pre-activated mutants may not be perfect surrogate of fully-active βarr conformations, the HDX pattern does suggest that the lariat loop region may adopt different conformations for activated βarr1 vs. βarr2, and it further complements our domain-swapping data described above.

Our functional data links the conformational differences between receptor-bound βarr1 and 2, as reported by Fab30-based sensor, with their different contributions in V_2_R desensitization. Previous studies have discovered that agonist-induced V_2_R endocytosis and ERK MAP kinase activation can be efficiently supported by the partially-engaged receptor-βarr complexes while receptor desensitization is driven primarily by the core-engaged complex (Cahill et al., 2017; Kumari et al., 2016; Kumari et al., 2017). This aligns with the possibility of differential core-engagement for βarr1 and 2 leading to their distinct contribution in receptor desensitization but comparable contribution in V_2_R endocytosis and ERK MAP kinase activation. It is also conceivable however that additional level of conformational differences in βarr isoforms may exist and drive the functional divergence with respect to endocytosis, signaling and ubiquitination for other GPCRs.

Our findings also raise some interesting questions and open new avenues for research in this area going forward. For example, some GPCRs such as muscarinic receptors contain a very short carboxyl-terminus but harbor phosphorylation sites in their 3^rd^ intracellular loops. Do such receptors also employ a biphasic mechanism of interaction with βarrs, and how do the two isoforms adopt distinct conformations for such receptors? Might there exist different conformations of receptor-bound βarr1 and 2 in response to stimulation by βarr-biased ligands and how do such conformational signatures govern the ensuing bias at the functional level? In addition, high-resolution structures of GPCR-βarr complexes, preferably of different βarr isoforms and different receptors, are still required to better understand the commonalities and differences in these signaling complexes. Future investigations to address some of these aspects should clearly offer novel insights into GPCR-βarr interaction and reveal how conformational differences in receptor-bound βarrs fine-tune their functional outcomes.

In conclusion, we discover structural and conformational differences between receptor-bound βarr isoforms which are potentially associated with their functional divergence in the context of GPCR regulatory and signaling paradigms. Our findings underline the importance of carefully considering both isoforms of βarrs when designing and characterizing βarr-biased GPCR ligands and thus, have direct implications for an ever-growing area of biased agonism aimed at designing novel GPCR therapeutics.

## ACKNOWLEDGEMENT

The research program in Dr. Shukla’s laboratory is supported by an Intermediate Fellowship of the Wellcome Trust/DBT India Alliance Fellowship [grant number IA/I/14/1/501285] awarded to AKS, the Science and Engineering Research Board (SERB) (EMR/2017/003804), Innovative Young Biotechnologist Award from the Department of Biotechnology (DBT) (BT/08/IYBA/2014-3) and the Indian Institute of Technology, Kanpur. Dr. Shukla is an Intermediate Fellow of Wellcome Trust/DBT India Alliance, EMBO Young Investigator and Joy Gill Chair Professor. Drs. Hemlata Dwivedi and Mithu Baidya were supported by National Post-Doctoral Fellowship of SERB (PDF/2016/002930 and PDF/2016/2893). Dr. Selent’s laboratory acknowledges support from the Instituto de Salud Carlos III FEDER (PI15/00460 and PI18/00094). JS and RGG also acknowledge the computer resources and technical support provided by the Barcelona Supercomputing Center (RES-BCV-2018-1-0012). TMS acknowledges support from National Center of Science, Poland (2013/08/M/ST6/00788 and 2017/27/N/NZ2/02571). Research in Dr. Chung’s laboratory is supported by the National Research Foundation of Korea funded by the Korean Government (NRF-2017K1A3A1A12072316). Dr. Dutta’s laboratory thanks DBT-IISc Partnership Program for EM consumables and transmission electron microscopy facility at Biological Sciences Division, IISc, Bangalore. Jaana Gastel, A Ph.D. student in the laboratory of Dr. Stuart Maudsley, is supported by the FWO-OP/Odysseus program (42/FA010100/32/6484) and her visit to the laboratory of Prof. Luttrell was funded by the FWO Travelling Fellowship Program (V4.161.17N). We thank Dr. Ramanuj Banerjee for his assistance with Figure 1 and Haaris Safdari for help in single particle negative-staining experiments. We are thankful to Prof. Ashwani K. Thakur for kindly allowing us to use the fluorometer in his laboratory for bimane experiments.

## AUTHORS’ CONTRIBUTION

Conceptualization, E.G., A.K.S. and J.S.; Methodology, E.G. A.K.S. and J.S.; Investigation, E.G., H.D., M.B., A.S., P.K., T.S., H.R.K., M.H.L., J.G., M.C., D.R., S.P., J.M., R.G.G. and S.D.; Writing – Original Draft, A.K.S.; Writing – Review & Editing, E.G., A.K.S. and J.S.; Funding Acquisition, A.K.S., L.M.L., K.Y.C., S.D. and J.S; Resources, L.M.L.; Supervision, A.K.S., L.M.L., K.Y.C., S.D. and J.S; Project Administration, A.K.S.

## STAR METHODS

### General reagents, construct design and protein expression

Commonly used reagents and cell culture consumables were purchased from Sigma-Aldrich or local vendors unless specified otherwise. *E. coli* expression constructs for βarr1 and 2, Fab30 and ScFv30 are described earlier, and these proteins were purified using previously described protocols (Kumari et al., 2017). βarr1^N245C^/βarr2^S246C^ were generated on a minimal cysteine background (i.e. harboring C59V, L68C, C125S, C140L, C150V, C242V, C251V, C269S mutations) by site-directed mutagenesis, and purified using a similar protocol as for wild-type. Expression constructs for β_2_V_2_R, GRK2, V_2_R, and their purification details have also been published previously (Kumari et al., 2017). Briefly, FLAG-β_2_V_2_R and FLAG-V_2_R were co-expressed with GRK2 in Sf9 cells (cultured in ESF921 media from Expression Systems), and 60-66h post-infection; cells were stimulated with indicated ligands and harvested by centrifugation.

### Sequence and structural analysis of βarrs

Phosphate interacting residues in V_2_Rpp-bound βarr1 were identified based on the previously determined crystal structure (PDB ID: 4JQI). They were compared to βarr2 by sequence alignment, structural visualization in PyMol and subsequent analysis in PDBsum as indicated in respective figure legends.

### ELISA assay

In order to assess the interaction of Fab30 with V_2_Rpp-bound βarrs (presented in Figure 1D and S3E), we first immobilized purified protein L (Genscript) onto MaxiSorp ELISA plates (Nunc). Subsequently, we incubated the wells with 1% BSA (Bovine Serum Albumin) to block non-specific binding. Afterwards, we added purified Fabs (in 20 mM Hepes, pH 7.4, 150mM NaCl; 1-3μg per well in 100μl volume) followed by gentle washing to remove unbound Fabs, and then added purified biotinylated βarrs (in 20 mM Hepes, pH 7.4, 150mM NaCl, 0.01% MNG; 1-3μg per well in 100μl volume) (with or without pre-incubation with V_2_Rpp). After an incubation of 15-30 min, wells were washed extensively (using 20mM Hepes, pH 7.4, 100mM NaCl, 0.01% MNG), and incubated with HRP-coupled streptavidin (Genscript; 1:2000 dilution of 1μg/ml). After another round of extensive washing, the reactivity of Fab30 with βarrs was visualized by adding TMB ELISA (Genscript). Colorimetric reaction was stopped by adding 2M H_2_SO_4_ and absorbance was measured at 450nm using a Victor X4 plate reader (Perkin-Elmer, USA).

In order to assess the recognition of receptor-bound βarrs by Fab30 (presented in S5A), we followed a recently described protocol (Kumari et al., 2016). Here, we first immobilized Fab30 on the ELISA plate followed by the addition of activated and phosphorylated receptor (in the form of cell lysate) mixed with purified βarrs. Their interaction was detected using HRP-coupled anti-FLAG M2 antibody (Sigma, 1:2,000 dilution) as the receptor contains an N-terminal FLAG tag.

### Co-immunoprecipitation assay

For coimmunoprecipitation based detection of Fab30 reactivity towards V_2_Rpp-bound βarrs (presented in Figure 1E, S3B-D and S8B), purified proteins were mixed together (in buffer containing 20mM Hepes, pH 7.4, 150mM NaCl, 0.01% MNG; 1:3 fold molar ratio of βarr:Fab; final concentrations in the range of 1-10μM) and incubated at room-temperature for 1h. Subsequently, protein L agarose beads were added to the reaction mix and incubated for an additional 1h at room-temperature. Beads were washed three times by centrifugation (using 20mM Hepes, pH 7.4, 150mM NaCl, 0.01% MNG), bound proteins were eluted using SDS sample buffer and separated by SDS-PAGE.

In order to evaluate the interaction of receptor-bound βarrs with Fabs by coIP, we used cell lysate prepared from S*f*9 cells co-expressing recombinant FLAG-tagged receptor and GRK2. Cells were first stimulated with an inverse agonist (to generate inactive and non-phosphorylated receptor) or an agonist (to generate activated and phosphorylated receptor). Cell lysate was pre-incubated with purified βarrs and Fab30 at room temperature for 1h. Subsequently, protein L beads (equilibrated in 20mM Hepes, pH 7.4, 100mM NaCl and 0.01% MNG) were added to the reaction mix and tumbled for an additional 1h. Beads were washed three times with washing buffer (same as equilibration buffer), bound proteins were eluted with SDS sample loading buffer and analyzed by Western blotting (HRP-coupled anti-FLAG M2 HRP from Sigma at 1:2000 dilution; HRP-coupled Protein L from Genscript at 1:2000 dilution).

In an alternative coIP set-up (presented in Figure S7D), HEK-293 cells expressing β_2_V_2_R, βarrmCherry and HA tagged ScFv30 as an intrabody, were stimulated with agonist or inverse agonist followed by coIP using anti-HA antibody coupled agarose beads (Sigma). Interaction of ScFv30 intrabody with βarr1 and 2 were visualized by Western blotting.

### Sortase-ligation protocol

For the preparation of receptor with homogenous phosphorylation, S*f*9 cells expressing β_2_AR (29-341) was first stimulated with 10nM BI-167107 for 30min and then resuspended in lysis buffer (50mM HEPES pH 7.4, 150mM NaCl, 10nM BI-167107, 1mM PMSF and 2mM benzamidine). Cells were lysed by dounce homogenization and lysate was solubilized in 1%(v/v) MNG for 2h at room temperature and cleared by centrifugation at 15000 rpm for 30min. Supernatant was incubated with pre-equilibrated M1-FLAG beads supplemented with 2mM CaCl_2_ for 2h at 4°C. Beads were washed alternately with low salt buffer(50mM HEPES pH 7.4, 150mM NaCl, 0.01% MNG, 10nM BI-167107 and 2mM CaCl_2_) and high salt buffer(50mM HEPES pH7.4, 350mM NaCl, 0.01% MNG, 10nM BI-167107 and 2mM CaCl_2_) respectively.

For ligation reaction, beads were resuspended in buffer containing 50mM Hepes, pH 7.4, 100mM NaCl, 0.01% (v/v) MNG, 10nM BI-167107, 5mM CaCl_2_, 50uM GGG-V2Rpp (GGGARGRpTPPpSLGPQDEpSCpTpTApSpSpSLAKDTSS) and 2 μM sortaseA. Slurry was incubated overnight at 4 °C and next day beads were alternately washed with low salt buffer and high salt buffer respectively. Ligated receptor was eluted with FLAG peptide and protein-L coIP assay was performed to assess the interaction of βarrs with receptor.

### Confocal microscopy

HEK-293 cells (ATCC) were cultured in DMEM (Sigma) supplemented with 5% Fetal Bovine Serum (Thermo Scientific) and 1% Penicillin-Streptomycin at 37°C under 5% CO_2_. Cells were transfected with indicated plasmids using PEI (Poly Ethylene Imine) at a DNA to PEI ratio of 1:3. 24h post-transfection, cells were split and seeded onto poly-L-lysine coated coverslips. After an additional 24h, cells were serum starved for 2h and then used for live cell imaging (LSM780NLO confocal microscope from Carl Zeiss) (Figure 2E and Figure S7B). Cells were stimulated with agonists for indicated time-points as mentioned in the respective figure legends.

### Bimane fluorescence assay

Experimental details of bimane labeling and fluorescence measurements have been described in detail previously (Kumari et al., 2016). Briefly, purified βarr1^N245C^ and βarr1^S245C^ were buffer exchanged in labeling buffer (20mM Hepes, 150mM NaCl, pH 7.5), and then incubated (approximately at a concentration of 1mg/ml) with freshly prepared monobromobimane (mBBr, Sigma-Aldrich) at a 10 fold molar excess. After 1h incubation on ice, unreacted mBBr was separated on a PD10 desalting column (GE Healthcare). Labeled βarrs were either used right away in the fluoresce measurements or they were flash frozen with 10% glycerol and stored at -80°C for later usage. For measuring the conformational change in the finger loop, mBBr labeled βarrs were mixed at 1:3 molar ratios with purified β_2_V_2_R. Purified receptors were also buffer exchanged in 20mM Hepes, pH 7.5, 150 mM NaCl, 0.01% MNG (Maltose Neopentyl Glycol), and final reactions were prepared such to maintain a consistent buffer (and detergent conditions). After incubating the mixture for 1h at room temperature, bimane fluorescence intensity was measured using a Fluorometer (Perkin Elmer, USA model LS-55) in photon counting mode as described previously (Kumari et al., 2016).

### Functional assays

HEK-293 cells were transfected with previously described and validated shRNAs targeting either βarr1 or 2 followed by generation of stable cells lines with puromycin selection using standard protocol (Vibhuti et al., 2011). Agonist-induced cAMP, receptor endocytosis and ERK MAP kinase phosphorylation was measured using previously described protocols (Kumari et al., 2017).

### Negative staining single particle analysis of β_2_V_2_R-βarr^swap2^-Fab30 complex

Samples for negative stain EM was prepared by conventional negative staining method (Ohi et al., 2004). Around 3.5 µl of the purified complex of β_2_V_2_-βarr^swap2^-Fab30 was adsorbed on glow discharged carbon coated copper grid for around one minute. This was followed by washing with three drops of water and staining with 0.5% uranyl formate for 30 seconds. Negatively stained β_2_V_2_-βarr^swap2^-Fab30 complex sample was imaged by using Tecnai T12 transmission electron microscope furnished with LaB6 filament and operating at 120kV accelerating voltage. Images were collected at magnification of 80k through side mounted Olympus VELITA (2K2K) CCD camera. All the images were collected at a defocus range of ∼-1.5 to -1.8µm yielding a final pixel size of ∼2 Å at specimen level. Total about 10000 particles from 150 micrographs were selected manually using e2boxer.py of EMAN 2.12 suite and used for 2D classification. The particle stack was classified into 50 classes using simple_prime2D script of SIMPLE software package.

### MD simulation set-up and analysis

#### Fab30 binding stability in solution

From the crystallized active βarr1 (PDB code: 4JQI), we removed the co-crystallized phosphopeptide and part of the Fab30 maintaining only residues 5 to 108 of the light chain and residues 1 to 123 of the heavy chain. The missing loop segment (309 to 310) of βarr1 was modeled using the loop modeler tool in the MOE software package (https://www.chemcomp.com/). Afterwards, we generated different inter-domain rotation states using linear interpolation between the active βarr1 (PDB code: 4JQI) and the inactive βarr1 (PDB code: 1G4R). Complexes were subjected to a geometrical optimization using the MOE package (CHARMM27 force field and born solvation). During this optimization, we applied constraints to the backbone atoms of the β-sheets and helices of βarr1.

To verify that obtained βarr1 conformations reflect a low-energetic conformation along the inactivation pathway, we compared interpolated structures to conformations observed in unbiased simulations of βarr1 inactivation (see Figure S9A). Interpolated structures were populated with an RMSD less than 1 Å in unbiased inactivation simulations proving low-energetic conformations.

Fab30 complex was obtained by the following procedure. To guarantee correct placement of Fab30 with respect to the C-domain (i.e. the main Fab30 binding interface) of the βarr1, all interpolation states were aligned to the C-domain of the active βarr1-Fab30 complex (PDB code: 4JQI). The Fab30 binding interface is optimized during the equilibration phase with only backbone atoms constrained (see section simulation setup below). Note that this procedure does not remove clashes of the Fab30 with the N-domain produced by low inter-domain rotation angles domain in the inactive βarr1 compared to the active state. In fact, these clashes are likely the reason for experimentally observed down regulation of Fab30 binding and are the focus of our simulation experiments.

The obtained βarr1-Fab30 complexes were solvated and ionized to 150 mM NaCl using VMD (Humphrey et al., 1996) yielding a system of approximately 86000 atoms. Systems were equilibrated for 10 ns in the NPT ensemble applying harmonic positional restraints to the protein backbone atoms and allowing side chains, water molecules and ions to relax (see detailed information below). Then, NVT production runs (see section simulation setup below) were used to assess Fab30 binding stability during simulation by monitoring the RMSD of the backbone atoms of the Fab30 β-sheet. In order to ensure a correct detection of Fab30 movement, we aligned the simulated complex to the C-domain of βarr1 prior to RMSD measurements.

#### Dynamics of the active and inactive βarr1

To sample the conformational flexibility of inactive and active βarr1 we started simulations from the active (PDB code: 4JQI) and inactive (PDB code: 1G4R) crystal structure. In the active structure, we removed the co-crystallized Fab30, maintaining the co-crystallized phosphopeptide to stabilize the active conformation. In both of the structures missing loops were modelled using the loop modeler tool in the MOE software package. Structures were solvated and ionized according to the protocol described above yielding systems above 56000 atoms. Afterwards, solvated complexes were subjected to equilibration and simulation (see section simulation setup below).

**Table 1.**
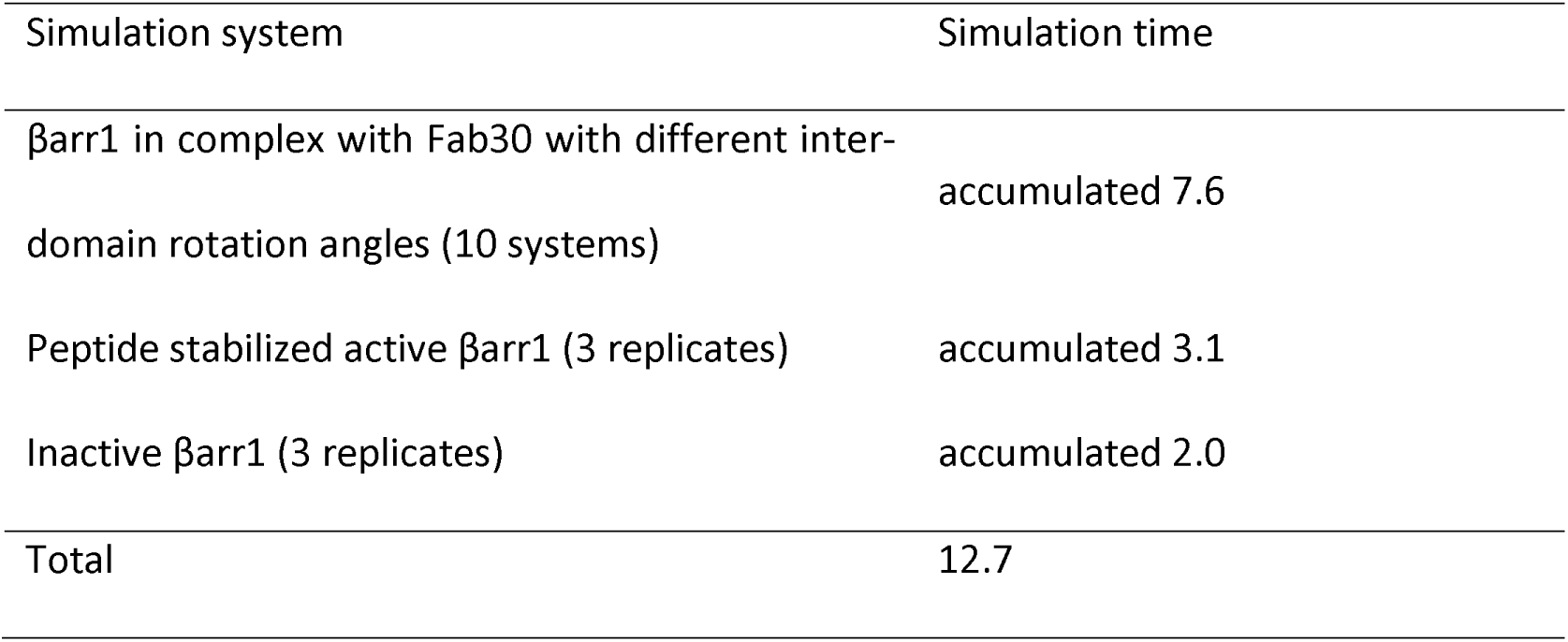
Simulation times (µs) for βarrs in receptor-bound complexes

#### Simulation setups

All simulations were carried out using the ACEMD simulation package (Harvey et al., 2009) and the CHARMM36m forcefield (Huang et al., 2017) and CHARMM36 forcefield (Klauda et al., 2010) force fields for proteins and lipids, respectively. NPT simulations were carried out at 310 K and 1 bar using the Berendsen barostat (H. J. Berendsen, 1984) with a relaxation time of 400 fs and 2 fs integration time step and harmonic constraints applied to all backbone atoms. NVT simulations were run at 310 K, using the Langevin thermostat (Grest and Kremer, 1986) with a damping coefficient of 5 ps^−1^ and 4 fs integration time step. No harmonic constraints are applied in this phase. In all simulations, we used a van der Waals and short-range electrostatic interactions with a cut-off of 9 Å and the particle mesh Ewald method (T. Darden, 1993) for long-range electrostatic interactions.

#### Analysis of inter-domain rotation angle

The inter-domain rotation angle is used as metric to assess the conformational landscape of βarrs in the receptor-bound state. For this purpose, we computed the displacement of the C-domain relative to the N-domain between the inactive (PDB code: 1G4R) and active βarr1 crystal structures (PDB code: 4JQI) as previously described in (Latorraca et al., 2018). The corresponding script was kindly provided by Naomi Latorraca.

#### Evaluation of inactivation pathway generated by interpolation

To verify that obtained βarr1 conformations reflect a low-energetic conformation along the inactivation pathway, we compared interpolated structures to conformations observed in unbiased simulations of βarr1 inactivation The active structure of βarr1 (PDB code: 4JQI) was solvated and simulated for 800ns in NVT conditions in three separate runs (see section simulation setup in main manuscript), allowing it to spontaneously inactivate.

Interpolation states were aligned to the frames of the βarr1 inactivation using the backbone atoms of β-sheets). Then, we quantified the presence of interpolated states along the inactivation pathway (Figure S9A). We find that all ten interpolated structures with an inter-domain rotation angle between 0° and 20° are sampled by unbiased simulation with an RMSD of β-sheets lower than 1. This indicates that generated states of βarr1 in Fab30 complexes adopt a low-energetic conformation along the inactivation pathway.

### Hydrogen/deuterium exchange analysis

Protein expression and purification conditions are as previously described (Yun et al., 2015). HDX-MS and data processing methods are also as previously described (Yun et al., 2015). Briefly, Purified protein was prepared in 60–100 µM in H_2_O buffer (20 mM HEPES, pH 7.4 and 150 mM NaCl). Hydrogen/deuterium exchange was initiated by mixing 2 µL of protein with 28 µL of D_2_O buffer (20 mM HEPES, pD 7.4, 150 mM NaCl in D_2_O), and the mixture was incubated for various time intervals (10, 100, 1000 and 10,000 s) on ice. At the indicated time points, the mixture was quenched by adding 30 µL of ice-cold quench buffer (100 mM NaH_2_PO_4_, pH 2.01). For non-deuterated (ND) samples, 2 µL of purified protein was mixed with 28 µL of H_2_O buffer to which, 30 µL of ice-cold quench buffer was added. Quenched samples were digested online by passing them through an immobilized pepsin column (2.1 × 30 mm) (Life Technologies, Carlsbad, CA, USA) at a flow rate of 100 µL/min with 0.05% formic acid in H_2_O at 11 °C. Peptide fragments were subsequently collected on a C18 VanGuard trap column (1.7 µm X 30 mm) (Waters, Milford, MA, USA) for desalting with 0.05% formic acid in H_2_O and were then separated by ultra-pressure liquid chromatography using an Acquity UPLC C18 column (1.7 µm, 1.0 X 100 mm) (Waters) at a flow rate of 35 µL/min with an acetonitrile gradient starting with 8% B and increasing to 85% B over 8.5 min. To minimize the back-exchange of deuterium to hydrogen, the system from trapping column to UPLC column was maintained at 0.5 °C and the buffers were adjusted to pH 2.5. Mass spectral analyses were performed with a Xevo G2 Qtof equipped with a standard ESI source (Waters). Mass spectra were acquired in the range of m/z 100–2000 for 12 min in positive ion mode. Peptide identification and HDX-MS data processing Peptic peptides were identified in non-deuterated samples with ProteinLynx Global Server 2.4 (Waters). Searches were run with the variable methionine oxidation modification. To process HDXMS data, the amount of deuterium in each peptide was determined by measuring the centroid of the isotopic distribution using the DynamX program (Waters). Back-exchange was not corrected because the data consisted of comparisons between β-arrestin1 and β-arrestin2 or between wild type and R169E mutants. All of the data was derived from at least three independent experiments.

### Receptor-βarr chemical cross-linking

For assessing receptor βarr interactions, Sf9 cells expressing β2V2R and GRK2 were first stimulated with an inverse agonist and agonist respectively for 30 minutes. Cells were lysed with lysis buffer (20mM Hepes pH 7.4, 100mM NaCl, 1X phosstop, 1mM PMSF and 2mM benzamidine) and incubated with purified βarr 1 or 2 at room temperature for 30 min. This was followed by the addition of 1mM dithiobis (succinimidyl-propionate) from a freshly prepared 100mM stock solution in DMSO. The lysate was incubated at room temperature for 40 min, and the reaction was quenched by adding 1M Tris pH 8.5. The lysate was solubilised in 1%(v/v) MNG for 1 h at room temperature and cleared by centrifugation at 15000 rpm for 30min. The supernatant was incubated with pre-equilibrated M1-FLAG beads supplemented with 2mM CaCl_2_ for 2h at 4ºC. Beads were washed alternately with low salt buffer(20mM Hepes pH 7.4, 150mM NaCl, 0.01% MNG, 2mM CaCl_2_) and high salt buffer(20mM Hepes pH7.4, 350mM NaCl, 0.01% MNG, 2mM CaCl2) respectively. Cross-linked proteins were eluted in FLAG-elution buffer (20mM Hepes pH 7.4, 150mM NaCl, 5mM EDTA, 0.01% MNG and 250ug/mL FLAG peptide). Samples were resolved on SDS-PAGE and visualized by Western blotting.

## Data analysis

Experimental data from ELISA assays, bimane fluorescence assay and densitometry based quantification of Western blots were plotted using GraphPad Prism software. The details of statistical analysis and number of biological replicates are indicated in the respective figure legends.

## Abbreviation

GPCRs: G Protein-Coupled Receptors;
βarrs: β-arrestins;
Fab: antigen binding fragment;
ScFv: single chain variable fragment;
β_2_AR: β2 adrenergic receptor;
V_2_R: vasopressin receptor subtype 2;
V_2_Rpp: V_2_R tail phosphopeptide;
mBBr: monobromobimane;
ERK MAP Kinase: extracellular signal regulated mitogen activated kinase;
ELISA: enzyme linked immunosorbent assay.

