## Supplemental Figures for "Conformational sensors and domain-swapping reveal structural and functional differences between β-arrestin isoforms"

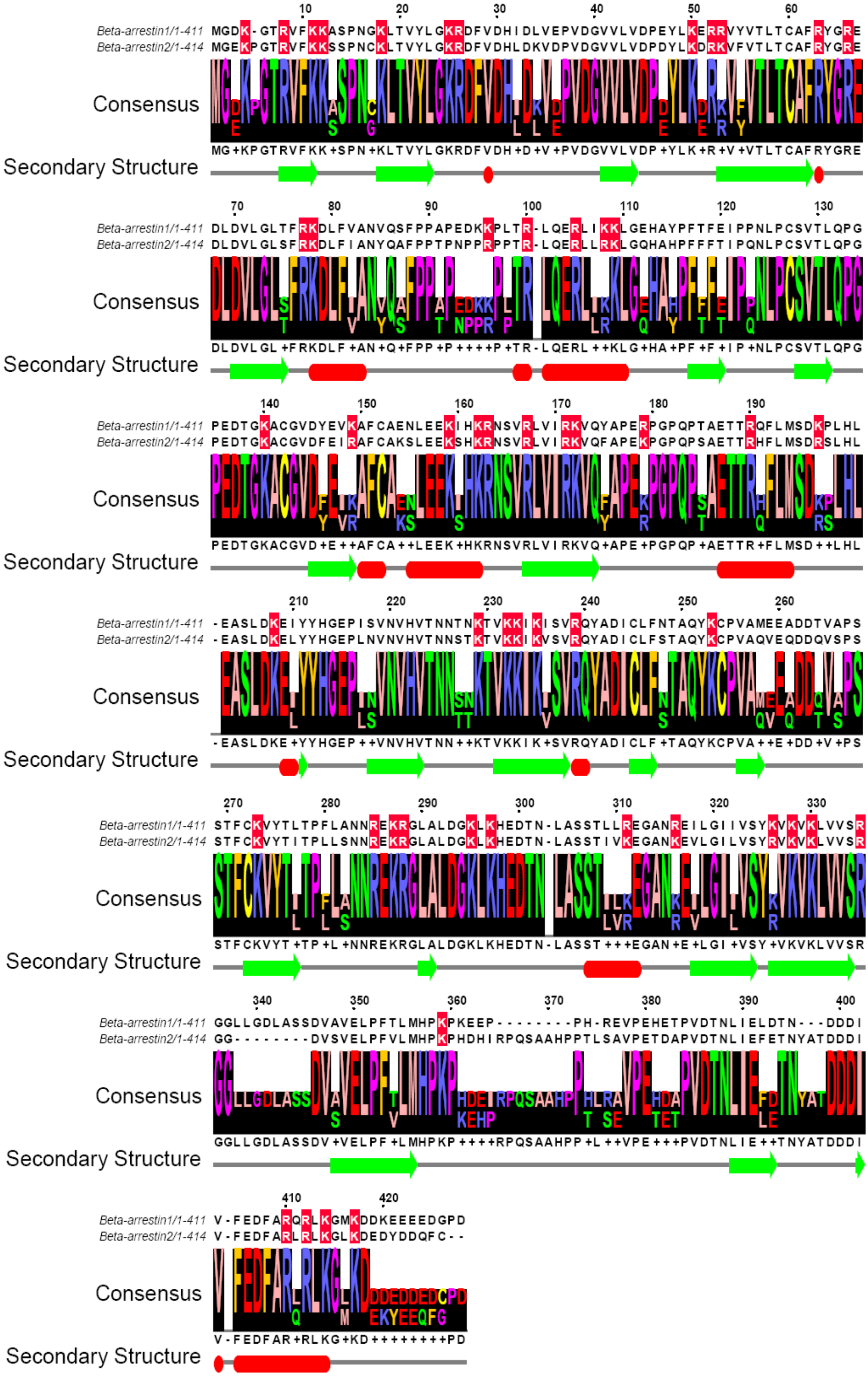


**Supplemental Figure 1. Overall sequence conservation between βarr1 and 2. Related to Figure 1.** Sequences of bovine βarr1 and 2 were aligned using the default T-Coffee and the alignment reliability was evaluated by Core/TCS tool. The aligned sequence was visualized and analyzed in Jalview program. Conserved Lys and Arg are highlighted in red. Shown below the alignment are Consensus Annotation and Sequence Logo. Secondary structure bar depicts JNetPRED annotation (The consensus prediction - helices are marked as red tubes, and sheets as dark green arrows) done by JNet secondary structure prediction in Jalview.

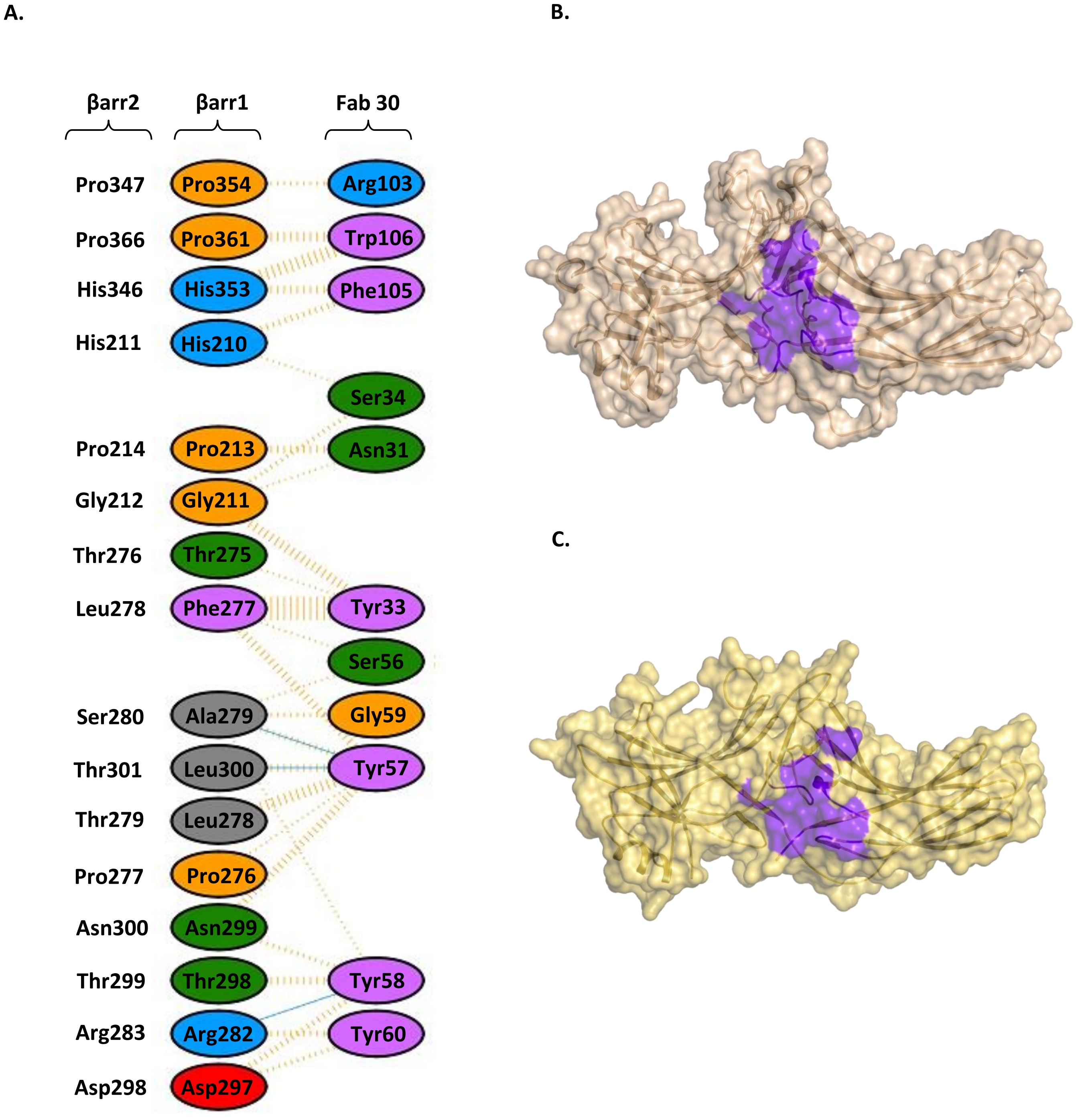


**Supplemental Figure S2. Fab30 interacting residues on βarr1 and 2 are well conserved. Related to Figure 2.** **A.** Schematic representation of amino acid residues in βarr1 that interact with Fab30 based on previously determined crystal structure (PDB ID: 4JQI). Coordinates of the crystal structure were submitted into *PDBSum,* and the interactions were mapped as a simplified ladder. **B.** The epitope of Fab30 on βarr1 is mapped on the crystal structure of $V_2$Rpp-bound βarr1. **C.** Amino acid residues in βarr1 that form the epitope of Fab30 are also conserved in βarr2 as mapped on the crystal structure (PDB ID: 3P2D).

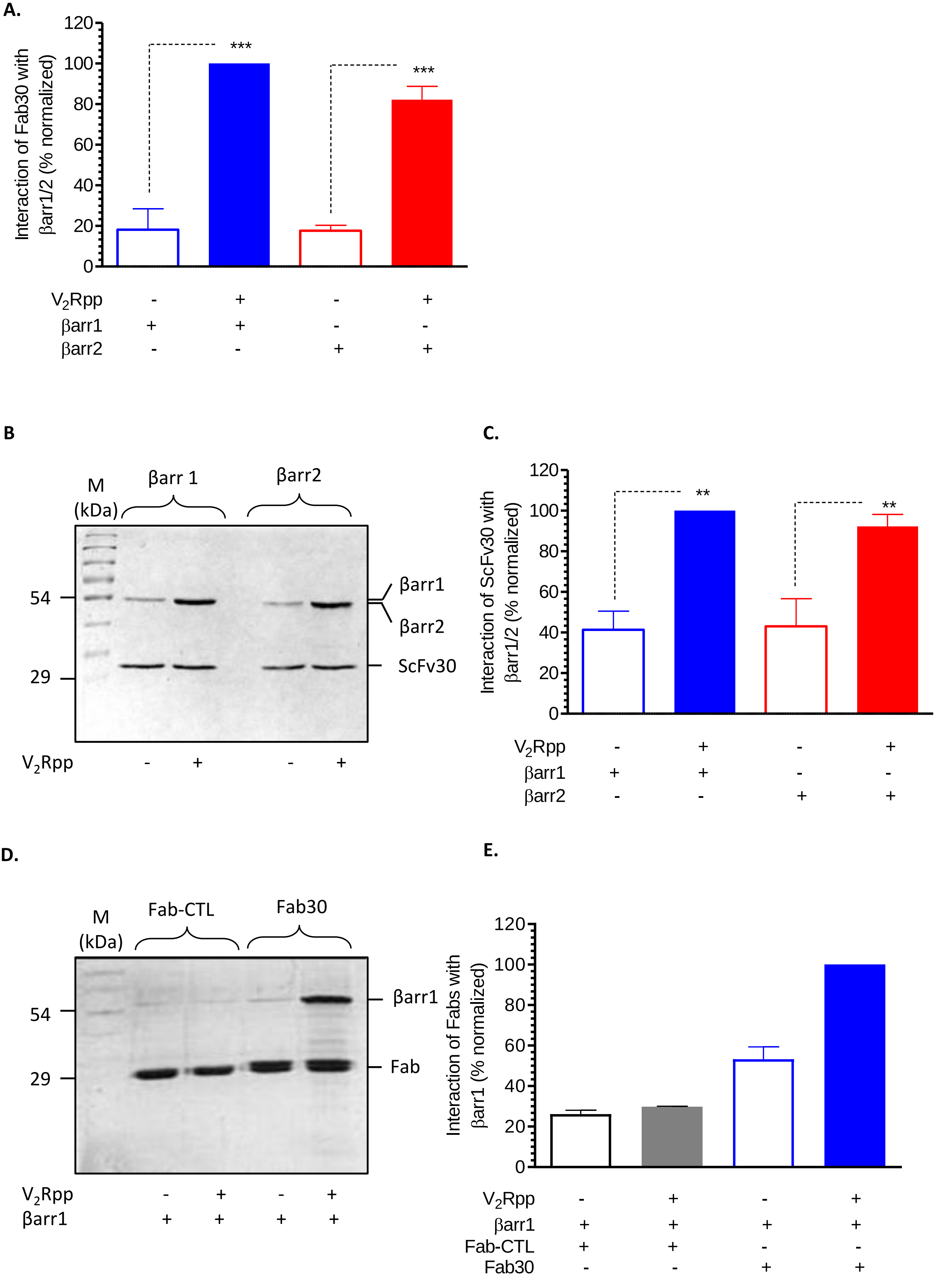
A.
Interaction of Fab30 with βarr1/2 (% normalized)
120
100
80
60
40
20
0
***
***
V2Rpp - + - +
βarr1 + + - -
βarr2 - - + +
B
M
(kDa)
βarr 1
βarr2
54
29
βarr1
βarr2
ScFv30
V2Rpp - + - +
C.
Interaction of ScFv30 with βarr1/2 (% normalized)
120
100
80
60
40
20
0
**
**
V2Rpp - + - +
βarr1 + + - -
βarr2 - - + +
D.
M
(kDa)
Fab-CTL
Fab30
54
29
βarr1
Fab
V2Rpp - + - +
βarr1 + + + +
E.
Interaction of Fabs with βarr1 (% normalized)
120
100
80
60
40
20
0
V2Rpp - + - +
βarr1 + + + +
Fab-CTL + + - -
Fab30 - - + +

**Supplemental Figure S3. Reactivity of Fab30 and ScFv30 towards $V_2$Rpp-bound βarr1/2. Related to Figure 2.** **A.** Densitometry-based quantification of data presented in Figure 1E. Average±SEM of three independent experiments are presented. **B.** Purified ScFv30 was incubated with $V_2$Rpp-bound βarr1 or 2 followed by co-immunoprecipitation using protein L beads. The interaction between ScFv30 and βarrs was visualized using Western blotting. **C.** Densitometry-based quantification of data presented in panel B. Average±SEM of three independent experiments are presented and data are normalized with respect to maximum signal for $V_2$Rpp-βarr1 condition (treated as 100%). Data were analyzed using One-Way ANOVA with Bonferroni post-test (***P<0.001; **P<0.01). **D-E.** A control Fab, referred to as Fab-CTL, that does not interact with βarrs, exhibits no significant reactivity towards $V_2$Rpp-bound βarr1 as assessed by co-IP assay. These experiments are carried out under the same experimental condition as in Figure 2A-B.

**A**

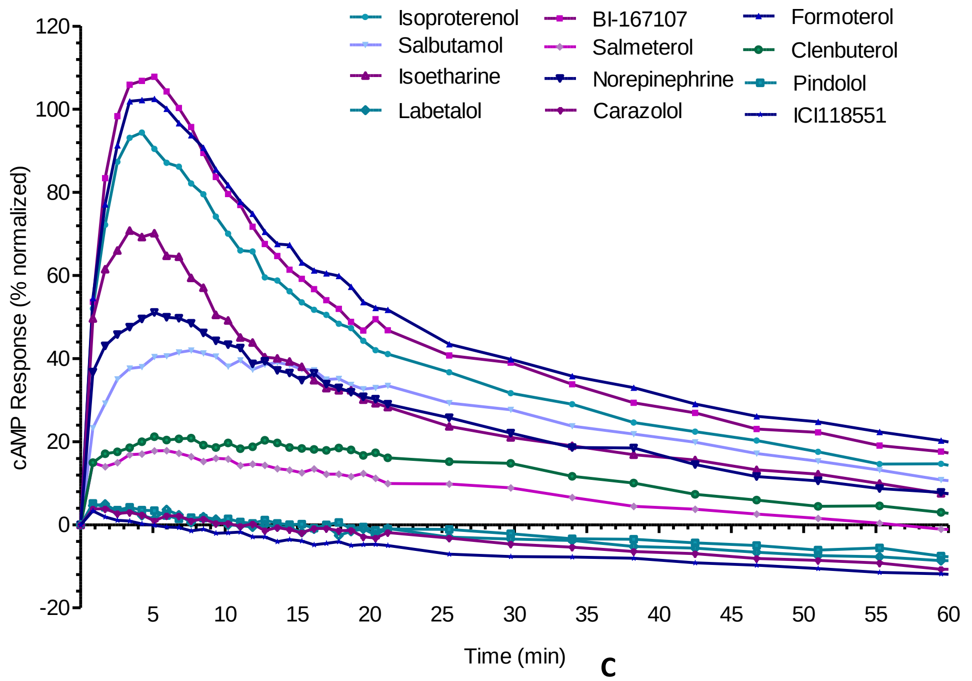


**B**

| Ligand | Integrated cAMP response |
|---|---|
| Isoproterenol | 100.0 ± 0.0 |
| BI-167107 | 122.6 ± 6.9 |
| Formoterol | 126.5 ± 8.6 |
| Salbutamol | 66.8 ± 9.0 |
| Salmeterol | 21.3 ± 5.1 |
| Clenbuterol | 29.8 ± 6.5 |
| Pindolol | 2.4 ± 1.4 |
| Isoetharine | 70.1 ± 9.1 |
| Norepinephrine | 63.8 ± 9.4 |
| Labetalol | 2.4 ± 1.5 |
| Cara | 2.0 ± 1.3 |
| ICI118551 | 1.0 ± 0.8 |

**C**

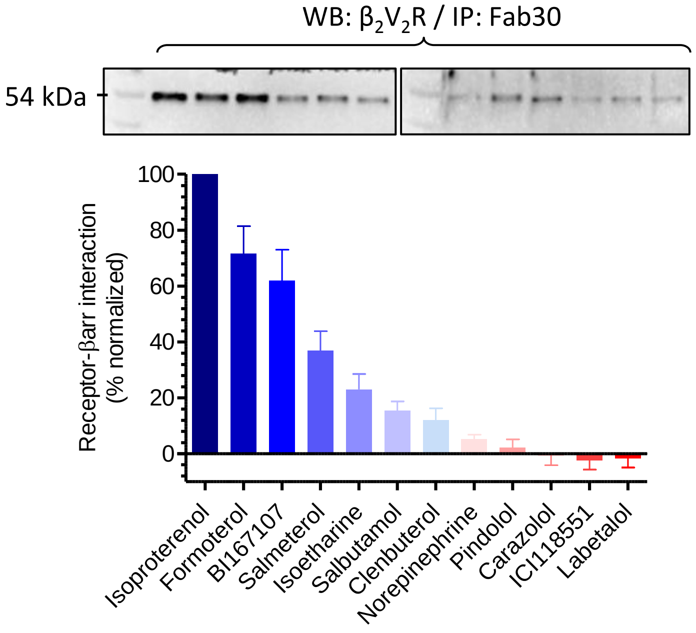


**D**

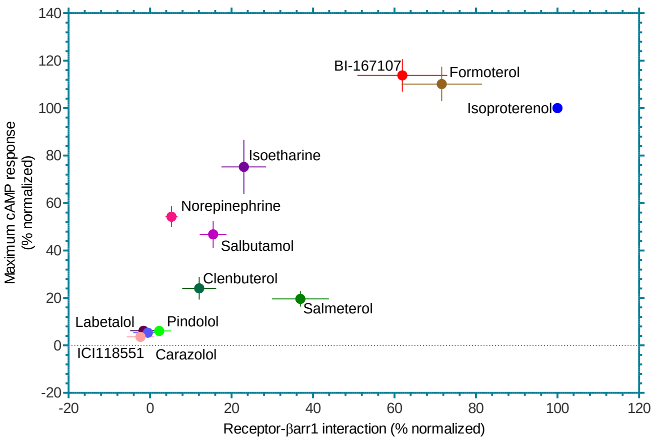


**E**

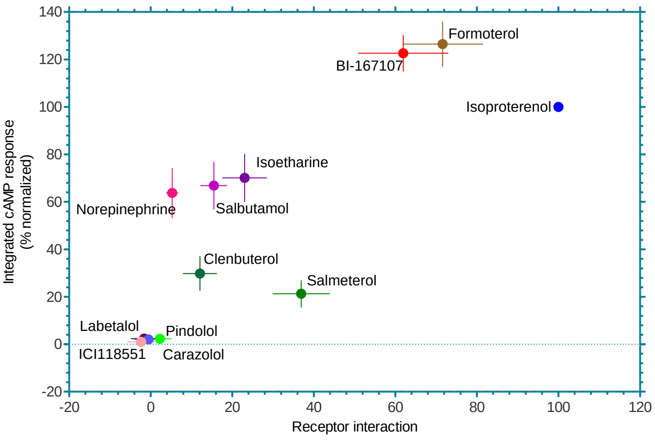

**Supplementary Figure S4. Correlation between ligand efficacy and Fab30 reactivity towards receptor-bound βarr1. Related to Figure 2. A.** cAMP response of twelve different $\beta_2$AR ligands using the Glo-sensor assay. HEK-293 cells expressing endogenous $\beta_2$AR were transfected with F-22 plasmid and stimulated with the saturating dose of ligands. Subsequently, the levels of cAMP were recorded for an extended time-period and plotted in GraphPad Prism. Data represent average of 5 independent experiments and normalized with isoproterenol response (treated as 100%). **B.** Quantitation of the integrated cAMP response (calculated as area under the curve) based on the data presented in panel A. The values in the right column indicated normalized response (±SEM) with respect to isoproterenol. **C.** Reactivity of Fab30 with receptor-bound βarr1. *Sf*9 cells expressing $\beta_2V_2$R were stimulated with saturating concentration of indicated ligands (1=Isoproterenol, 2=BI167107, 3=Formoterol,4=Salbutamol, 5=Salmeterol, 6=Clenbuterol, 7=Pindolol, 8=Isoetharine, 9=Norepinephrine, 10=Labetalol, 11=Carazolol, 12=ICI118551). Subsequently, cells were lysed, incubated with purified βarr1 and Fab30 followed by co-immunoprecipitation using protein L beads. Samples were subsequently visualized by Western blotting using HRP-coupled anti-FLAG M2 antibody. A representative image from six independent experiments is presented here and the densitometry-based quantification of all six experiments is presented in the lower panel. **D-E.** Correlation between cAMP response (maximum values or integrated response based on data presented in panel A-B) and the reactivity of Fab30 towards receptor-bound βarr1 as presented in panel C.

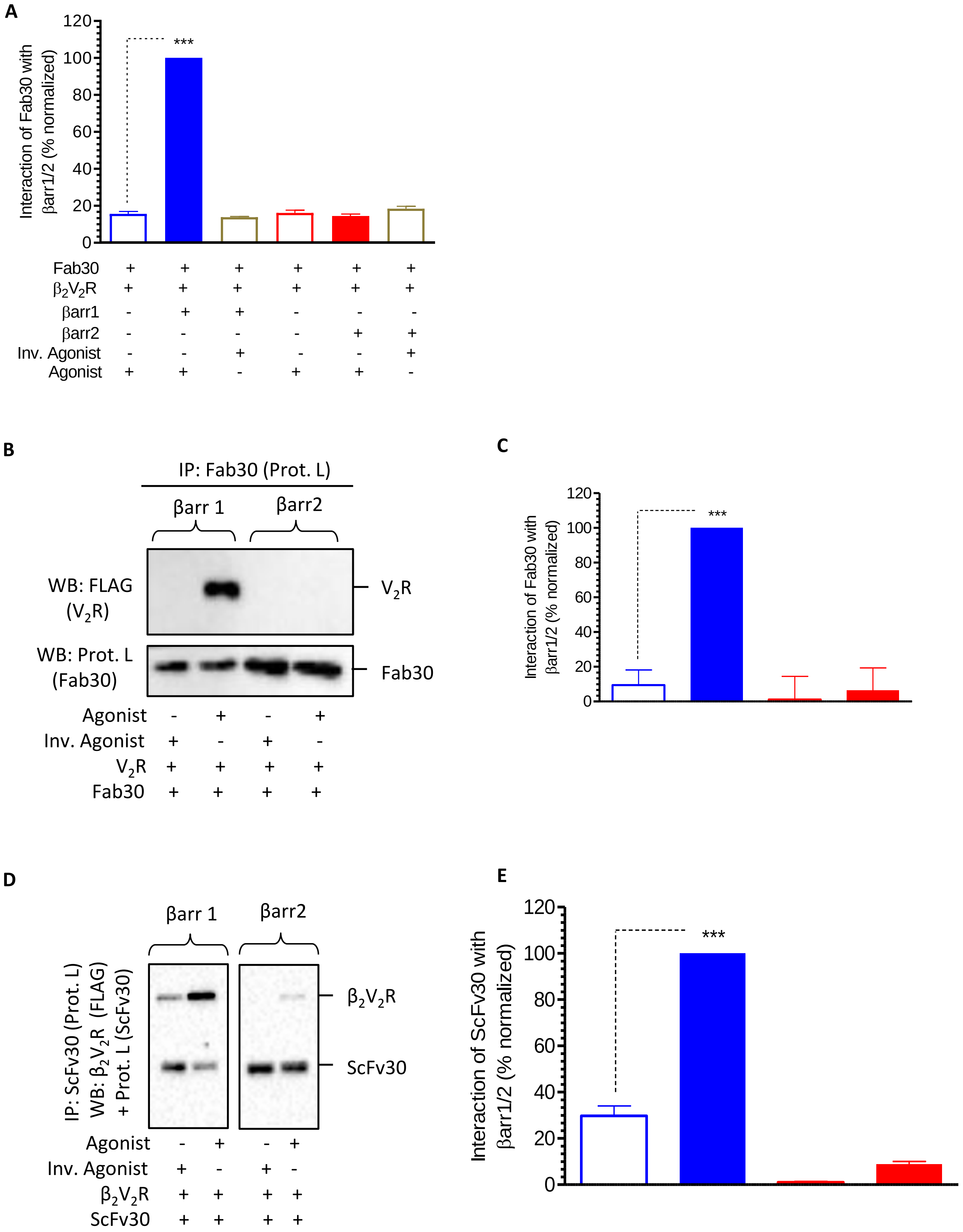

A
Interaction of Fab30 with βarr1/2 (% normalized)
120
100
80
60
40
20
0
***
Fab30 + + + + + +
β2V2R + + + + + +
βarr1 - + + - - -
βarr2 - - - - + +
Inv. Agonist - - + - - +
Agonist + + - + + -
B
IP: Fab30 (Prot. L)
βarr 1
βarr2
WB: FLAG (V2R)
V2R
WB: Prot. L (Fab30)
Fab30
Agonist - + - +
Inv. Agonist + - + -
V2R + + + +
Fab30 + + + +
C
Interaction of Fab30 with βarr1/2 (% normalized)
120
100
80
60
40
20
0
***
D
βarr 1
βarr2
IP: ScFv30 (Prot. L)
WB: β2V2R (FLAG) + Prot. L (ScFv30)
β2V2R
ScFv30
Agonist - + - +
Inv. Agonist + - + -
β2V2R + + + +
ScFv30 + + + +
E
Interaction of ScFv30 with βarr1/2 (% normalized)
120
100
80
60
40
20
0
***

**Supplemental Figure S5. Reactivity of Fab30/ScFv30 towards receptor-bound βarr1 and 2. Related to Figure 2. A.** The reactivity of Fab30 towards $\beta_2V_2R$-bound βarr1 and 2 were measured in an ELISA assay. First, purified Fab30 was immobilized in polystyrene plates, followed by the addition of $\beta_2V_2R$ and either βarr1 or 2. Afterwards, the wells were washed extensively and the captured $\beta_2V_2R$ was visualized using HRP-coupled anti-FLAG M2 antibody. Data represent average±SEM of three independent experiments, each carried out in duplicate. Data are normalized with respect to maximum signal for agonist+$\beta_2V_2R$+βarr1_Fab30 condition (treated as 100%) and analyzed using One-Way ANOVA with Bonferroni post-test (***$P<0.001$). **B-C.** Co-immunoprecipitation assay reveals selective recognition of $V_2R$-bound βarr1 but not βarr2 by Fab30. This coIP experiment was performed in a similar fashion as described in Figure 2A. Panel C shows densitometry based quantification of data, and values represent mean±SEM of three independent experiments analyzed using One-Way ANOVA with Bonferroni post-test (***$P<0.001$). Data are normalized with respect to agonist-$V_2R$-βarr1**-**Fab30 condition (treated as 100%). Here, Tolvaptan (1µM) is used as an inverse agonist while AVP (1µM) is used as an agonist to stimulate the cells expressing FLAG-tagged $V_2R$. **D-E.** Co-immunoprecipitation assay reveals selective recognition of receptor-bound βarr1 but not of βarr2 by ScFv30. This coIP experiment was performed in a similar fashion as described for Fab30 in Figure 2A. Panel E shows densitometry based quantification of data, and values represent mean±SEM of three independent experiments analyzed using One-Way ANOVA with Bonferroni post-test (***$P<0.001$). Data are normalized with respect to agonist-$\beta_2V_2R$-βarr1-ScFv30 condition (treated as 100%).

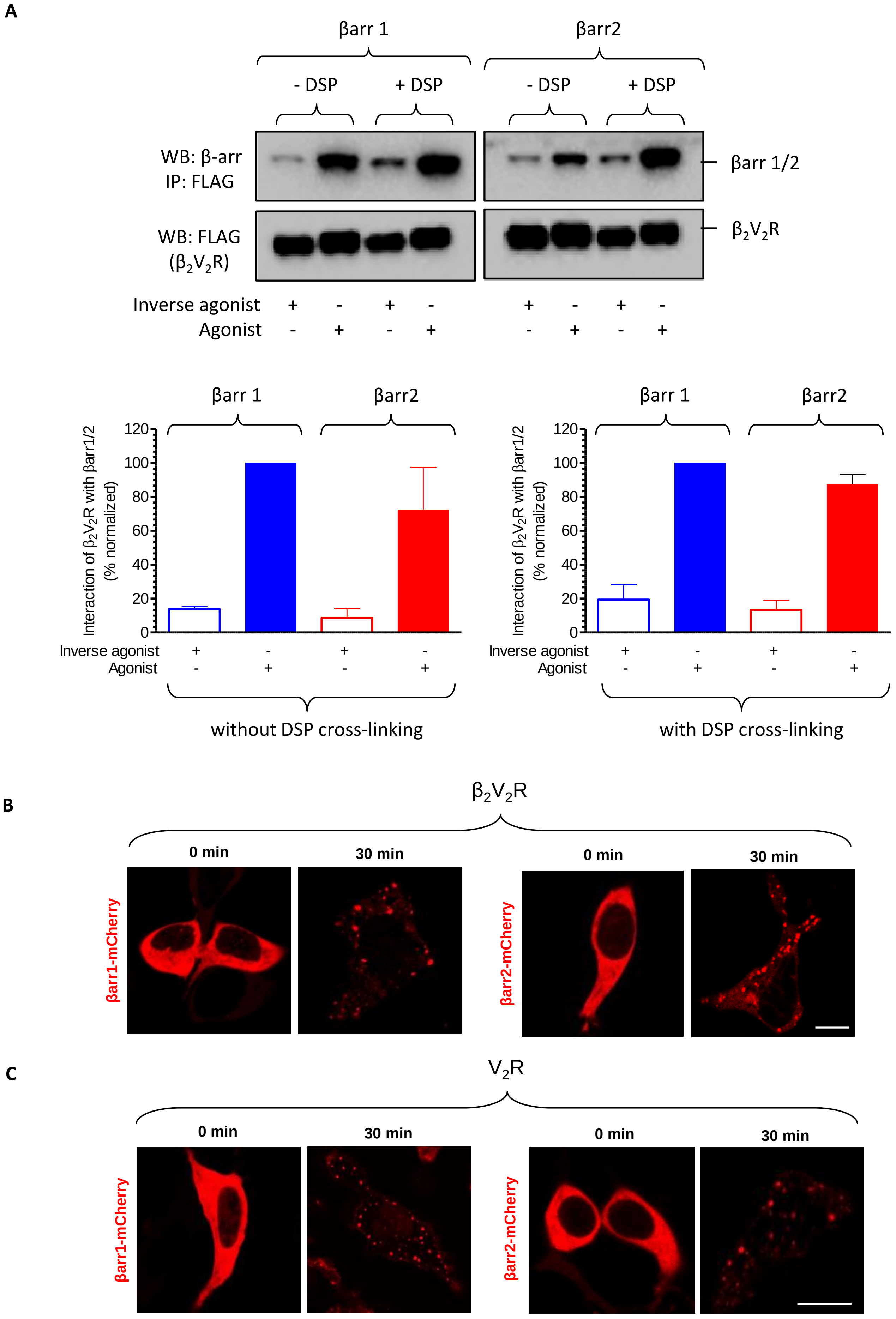
A
βarr 1
βarr2
- DSP
+ DSP
- DSP
+ DSP
WB: β-arr
IP: FLAG
WB: FLAG
(β2V2R)
βarr 1/2
β2V2R
Inverse agonist + - + - + - + -
Agonist - + - + - + - +
βarr 1
βarr2
Interaction of β2V2R with βarr1/2 (% normalized)
0
20
40
60
80
100
120
Inverse agonist + - + -
Agonist - + - +
without DSP cross-linking
with DSP cross-linking
B
β2V2R
0 min
30 min
βarr1-mCherry
βarr2-mCherry
C
V2R
0 min
30 min
βarr1-mCherry
βarr2-mCherry

**Supplemental Figure S6. $\beta_2V_2R$ and $V_2R$ recruits βarr2 upon agonist-stimulation. Related to Figure 2.** *Sf*9 cells expressing $\beta_2V_2R$ were stimulated with either an inverse agonist (carazolol) or full agonist (BI-167107). Subsequently, cells were lysed and equal amounts of purified βarr 1 or 2 were added. Subsequently, $\beta_2V_2R$ was immunoprecipitated using anti-FLAG M2 antibody agarose and the interaction of βarr 1 and 2 was visualized by Western blotting. In another set, a cross-linking reaction was also performed with 1mM DSP (dithiobis succinimidyl propionate) after mixing the cell lysate with βarrs followed by coimmunoprecipitation experiment. We observed comparable interaction of both, βarr1 and 2 with $\beta_2V_2R$ which confirms that the lack of reactivity of Fab30 with receptor-bound βarr2 is not because of the inability of the receptor to interact with βarr2 or due to the biochemical quality of purified βarr2 protein. These experiments were carried out twice with identical results and a representative image is shown. The lower panels show densitometry based quantification of $\beta_2V_2R$-βarr interaction, and data are normalized with respect to agonist-$\beta_2V_2R$-βarr1 condition (treated as 100%). **B.** $\beta_2V_2R$ recruits βarr2 upon agonist-stimulation as visualized by confocal microscopy. HEK-293 cells expressing $\beta_2V_2R$ and βarr1/2-mCherry were stimulated with 10µM isoproterenol for 30 min. Subsequently, the recruitment and internalization of $\beta_2V_2R$-bound βarr1/2 in endosomal vesicles was visualized by confocal microscopy. Scale bar is 10µm. **C.** $V_2R$ recruits βarr1/2 upon agonist-stimulation as visualized by confocal microscopy. HEK-293 cells expressing $V_2R$ and βarr-mCherry were stimulated with either AVP (100nM) for 30 minutes. Subsequently, the recruitment and internalization of $V_2R$-bound βarr2 in endosomal vesicles was visualized by confocal microscopy. Scale bar is 10µm.

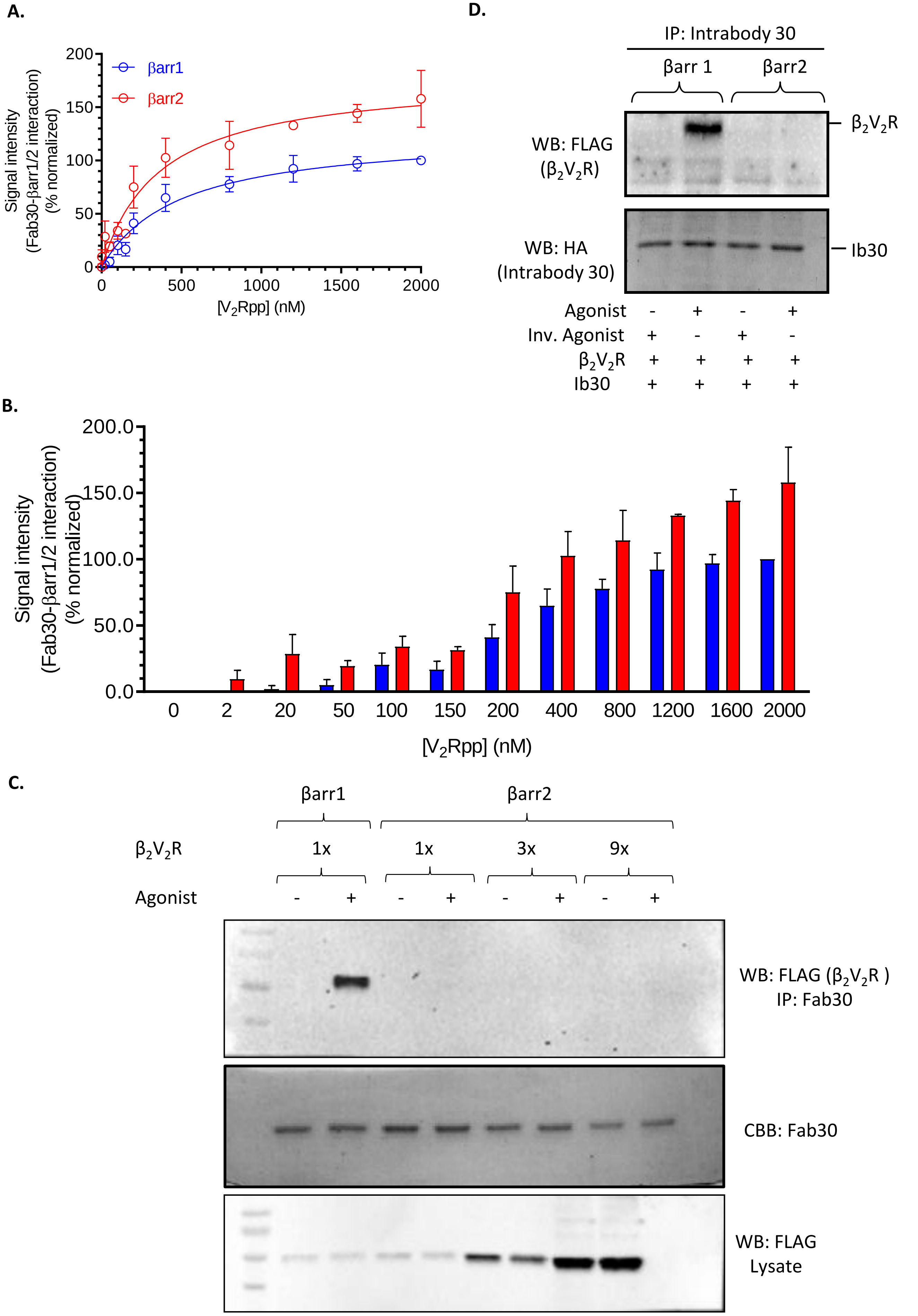
A.
βarr1
βarr2
Signal intensity
(Fab30-βarr1/2 interaction)
(% normalized)
0
50
100
150
200
0
500
1000
1500
2000
[V2Rpp] (nM)
B.
Signal intensity
(Fab30-βarr1/2 interaction)
(% normalized)
0.0
50.0
100.0
150.0
200.0
0
2
20
50
100
150
200
400
800
1200
1600
2000
[V2Rpp] (nM)
C.
βarr1
βarr2
β2V2R
1x
1x
3x
9x
Agonist
- + - + - + - +
WB: FLAG (β2V2R )
IP: Fab30
CBB: Fab30
WB: FLAG
Lysate
D.
IP: Intrabody 30
βarr 1
βarr2
WB: FLAG
(β2V2R)
β2V2R
WB: HA
(Intrabody 30)
Ib30
Agonist - + - +
Inv. Agonist + - + -
β2V2R + + + +
Ib30 + + + +

**Supplemental Figure S7. Reactivity of Fab30 towards $V_2$Rpp-bound and receptor-bound βarr1 and 2. Related to Figure 2. (A-B)** Purified βarr1/2 (200nM) were incubated with increasing concentrations of $V_2$Rpp (as indicated in the figures) followed by ELISA experiment to measure the reactivity by Fab30. We observed that Fab30 recognizes $V_2$Rpp-bound βarr2 as efficiently as βarr1, even at partial occupancy with $V_2$Rpp. This observation suggest that there is no major affinity difference of Fab30 for $V_2$Rpp-bound βarr1 and 2. Data represent average±SEM of three independent experiments, each carried out in duplicate. **(C)** Purified βarr1/2 were incubated with Fab30 and cellular lysate prepared from *Sf*9 cells expressing $\beta_2V_2R$. Despite using up-to nine-fold more $\beta_2V_2R$, we still did not observe any detectable reactivity of Fab30 towards receptor-bound βarr2. This finding indicates that the lack of Fab30 reactivity is not due to stoichiometric differences in terms of available phosphorylated carboxyl-terminus. **(D)** Intrabody 30 (Ib30) confirms distinct conformations of receptor-bound βarr1 and 2 in cellular context. HEK-293 cells were transfected with $\beta_2V_2R$, βarr1/2 and Ib30 followed by co-immunoprecipitation experiment using the HA tag on Ib30. The interaction of Ib30 with receptor-βarr complexes was visualized by Western blotting. A representative image from two independent experiments is presented here.

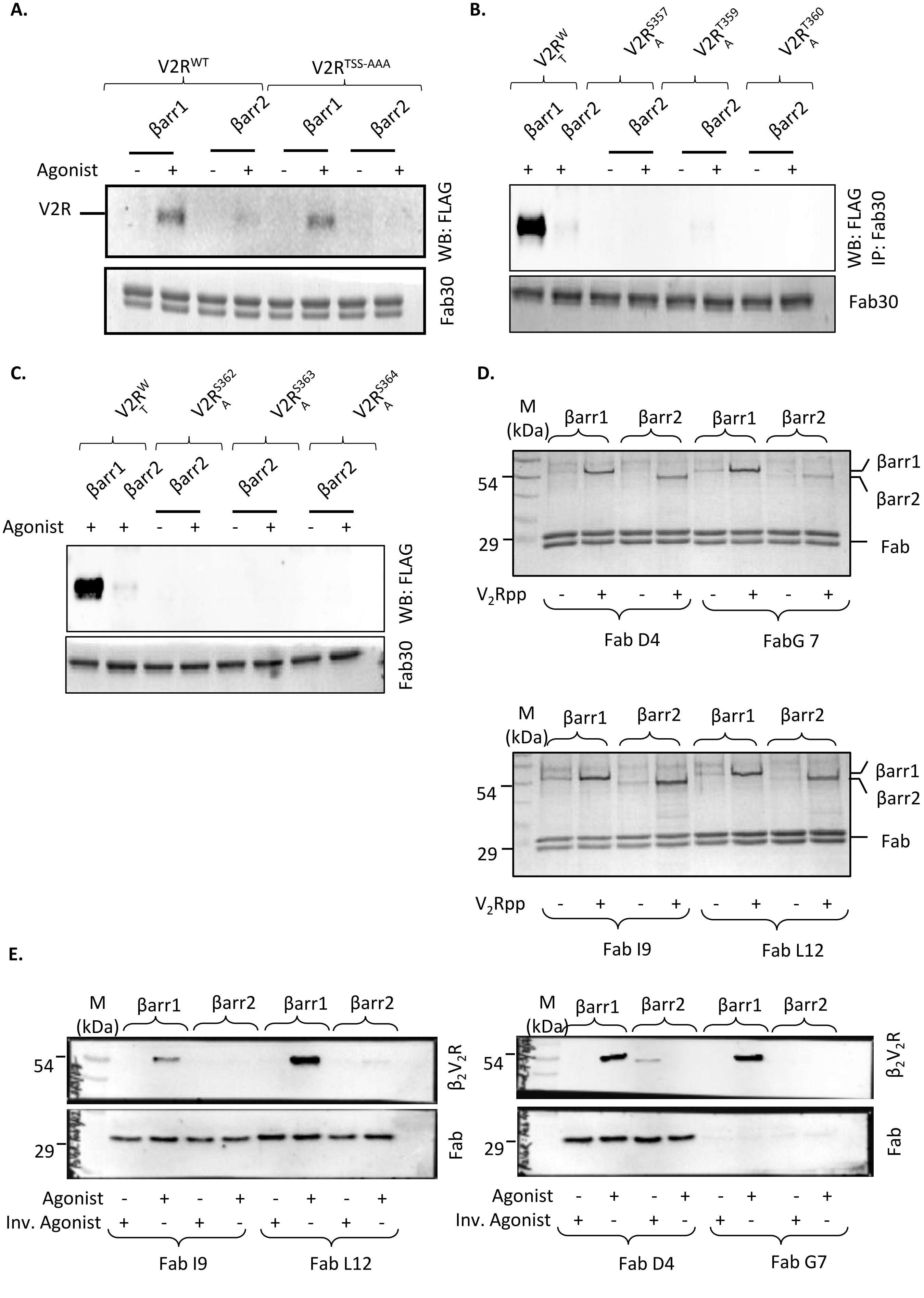
A.
V2R^WT
V2R^TSS-AAA
βarr1
βarr2
βarr1
βarr2
Agonist - + - + - + - +
V2R
WB: FLAG
Fab30
B.
V2R^WT
V2R^S357A
V2R^T359A
V2R^T360A
βarr1
βarr2
βarr2
βarr2
βarr2
+ + - + - + - +
WB: FLAG
IP: Fab30
Fab30
C.
V2R^WT
V2R^S362A
V2R^S363A
V2R^S364A
βarr1
βarr2
βarr2
βarr2
βarr2
Agonist + + - + - + - +
WB: FLAG
Fab30
D.
M (kDa)
βarr1
βarr2
βarr1
βarr2
54
29
βarr1
βarr2
Fab
V2Rpp - + - + - + - +
Fab D4
FabG 7
M (kDa)
βarr1
βarr2
βarr1
βarr2
54
29
βarr1
βarr2
Fab
V2Rpp - + - + - + - +
Fab I9
Fab L12
E.
M (kDa)
βarr1
βarr2
βarr1
βarr2
54
29
β2V2R
Fab
Agonist - + - + - + - +
Inv. Agonist + - + - + - + -
Fab I9
Fab L12
M (kDa)
βarr1
βarr2
βarr1
βarr2
54
29
β2V2R
Fab
Agonist - + - + - + - +
Inv. Agonist + - + - + - + -
Fab D4
Fab G7

**Supplemental Figure S8. Site-directed mutagenesis and additional Fab sensors corroborate the structural differences between βarr1 and 2. Related to Figure 2.** (**A-C)** A series of $V_2R$ mutants were generated harboring site-specific mutations of distinct phosphorylation sites in the carboxyl-terminus. HEK-293 cells expressing either WT or mutant $V_2R$ constructs were stimulated with agonist (AVP; 100nM, 30 min), lysed, incubated with purified βarr1/2 and Fab30. Subsequently, co-IP experiment was performed using protein L agarose beads and proteins were detected using Western blot. Similar to WT receptor, Fab30 did not exhibit any detectable recognition of receptor-bound βarr2 for any of the receptor mutants. A representative image of two independent experiments are presented here. **(D)** A set of additional Fabs reveal conformational similarity between $V_2Rpp$-bound βarr1 and 2. Purified βarr1 or 2 were incubated with the indicated Fab in absence or presence of $V_2Rpp$ followed by co-immunoprecipitation using Protein L beads and visualization using SimplyBlue staining. A representative image of two independent experiments is shown. **(E)** Additional Fabs reveal distinct conformations of $\beta_2V_2R$-bound βarr1 and 2. *Sf*9 cells expressing FLAG-$\beta_2V_2R$ were stimulated with either an agonist (BI167107, 1μM) or inverse agonist (Carazolol, 1μM) for 1h followed by incubation with purified βarr1 or 2 and indicated Fabs. Afterwards, the receptor was solubilized using 1% MNG and co-immunoprecipitated using Protein L beads. Samples were visualized by Western blotting using HRP-coupled anti-FLAG M2 antibody (for $\beta_2V_2R$) and HRP-coupled Protein L (for Fabs). For some reason, FabG7 is not stained well with HRP-coupled Protein L but Ponceau staining of the membrane shows equal amount of Fab pull-down.

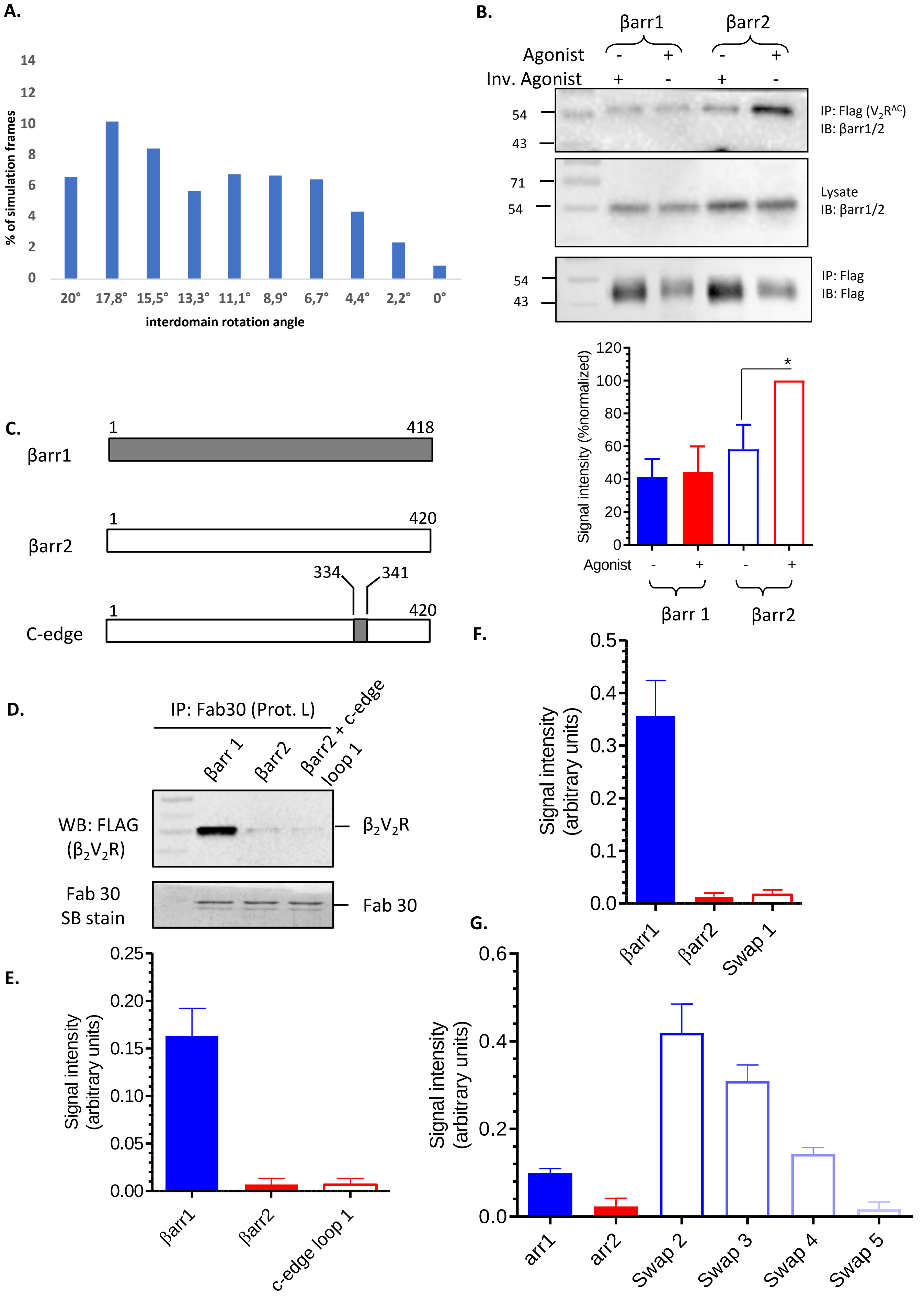

A.
% of simulation frames
interdomain rotation angle
20° 17,8° 15,5° 13,3° 11,1° 8,9° 6,7° 4,4° 2,2° 0°
B.
βarr1
βarr2
Agonist
Inv. Agonist
IP: Flag ($V_2R^{\Delta C}$)
IB: βarr1/2
Lysate
IB: βarr1/2
IP: Flag
IB: Flag
Signal intensity (%normalized)
Agonist
βarr 1
βarr2
C.
βarr1
βarr2
C-edge
D.
IP: Fab30 (Prot. L)
Βarr 1
Βarr2
Βarr2 + c-edge loop 1
WB: FLAG ($\beta_2V_2R$)
$\beta_2V_2R$
Fab 30 SB stain
Fab 30
E.
Signal intensity (arbitrary units)
Βarr1
Βarr2
c-edge loop 1
F.
Signal intensity (arbitrary units)
Βarr1
Βarr2
Swap 1
G.
Signal intensity (arbitrary units)
arr1
arr2
Swap 2
Swap 3
Swap 4
Swap 5

**Supplemental Figure S9. Population of interpolated states within an inter-domain rotation angle (0° to 20°) in unbiased simulations and Reactivity of Fab30 with chimeric βarrs. Related to Figures 4-5. (A)** The presence of interpolated states along the inactivation pathway (unbiased simulation) is quantified as percentage of frames with an RMSD of backbone atoms of β-sheets < 1 Å. . **(B)** HEK-293 cells were transfected with carboxyl-terminus truncated $V_2R$ i.e. Flag-$V_2$RC-term and βarr1/2 followed by stimulation with indicated ligands, cross-linking and co-immunoprecipitation using anti-Flag antibody agarose. Proteins were visualized on Western blots using corresponding antibodies. A representative image of three independent experiments is shown. The lower panel shows densitometry-based quantification of data is presented after normalization with respect to agonist+βarr2 condition treated as 100%. **(C)** Schematic representation of βarr2 construct with grafted C-edge loop 1 from βarr1. **(D)** Co-immunoprecipitation experiment reveals that Fab30 fails to detect receptor-bound conformation of βarr2 construct with grafted C-edge loop 1. This co-IP experiment was performed following the same protocol as described in Figure 2A. This data suggest that membrane anchoring function of C-edge loop 1, and the lack of such a possibility in βarr2 is not responsible for the Fab30 reactivity pattern. A representative image of three independent experiment is presented here. **(E)** Densitometry based quantification of three independent experiments. **(F)** Densitometry based quantification of data presented in Figure 4B showing the Fab30 reactivity patterns towards βarr1, 2 and swap1. **(G)** Densitometry based quantification of data presented in Figure 4D showing the Fab30 reactivity patterns towards βarr1, 2 and swap2-5.

**A.**

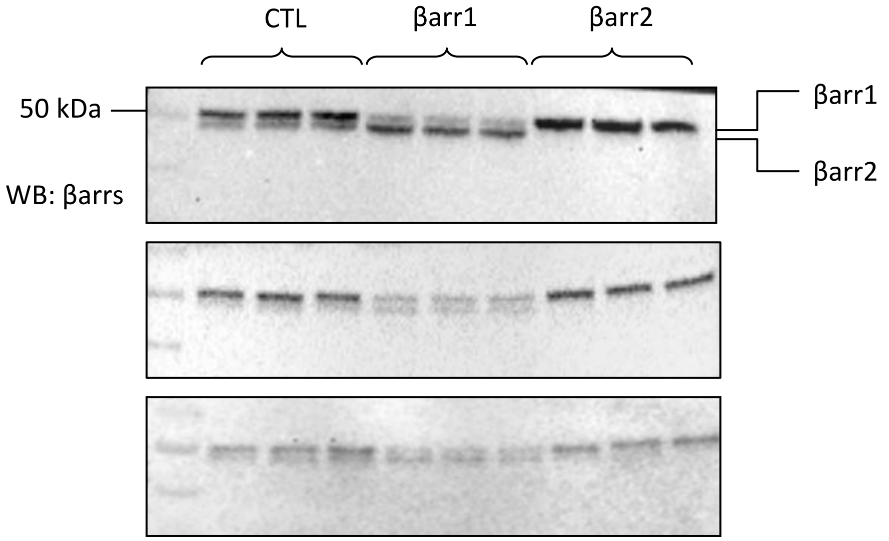


**B.**

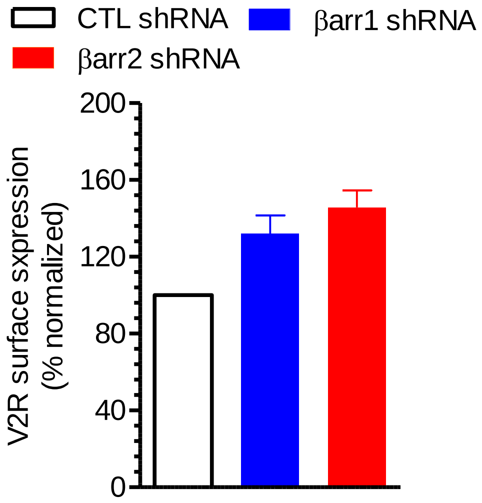


**C.**

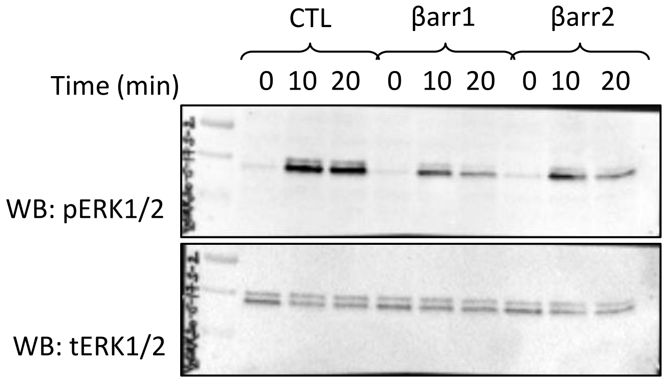


**Supplemental Figure S10. βarr knock-down, receptor surface expression and ERK1/2 phosphorylation. Related to Figure 5. (A)** Western blot analysis of lysates prepared from HEK-293 cells stably expressing CTL-shRNA, βarr1-shRNA and βarr2-shRNA to measure βarr knock-down levels**.** Lysates from three different wells of a six-well plate are shown from three different experiments. **(B)** Surface expression of $V_2R$ was comparable in cells transfected with CTL, βarr1 and βarr2 shRNA as measured by whole cell ELISA assay. Data are normalized with respect to $V_2R$ expression in CTL shRNA transfected cells (treated as 100%). **(C)** Agonist-induced ERK activation downstream of $V_2R$ depends on both βarr1 and 2 as revealed by comparing the ERK phosphorylation in cells transfected with CTL, βarr1 and βarr2 shRNA. These experiments were carried out five times with identical results and a representative image is shown. Densitometry based quantification of data from all five experiments presented in Figure 5D.

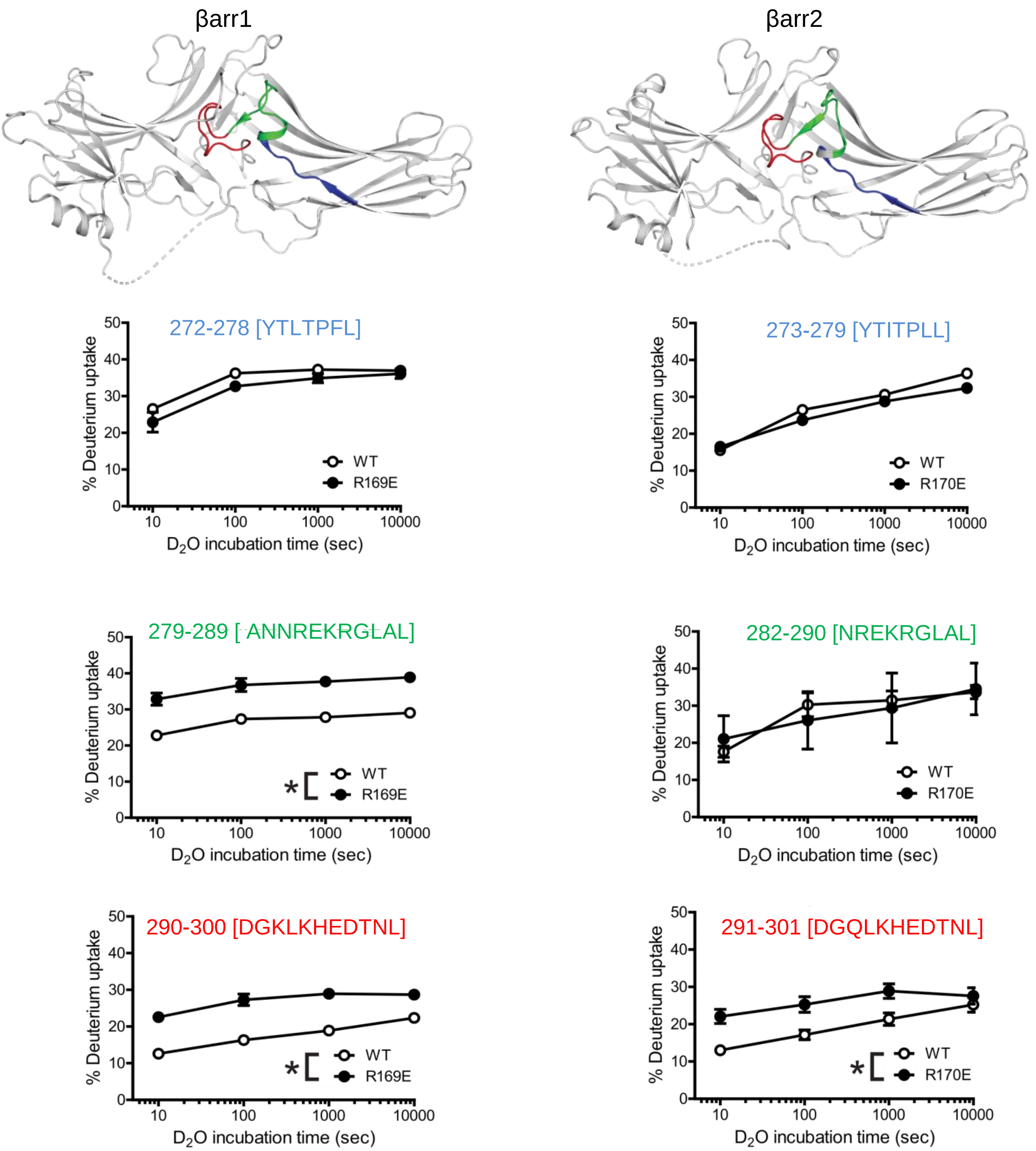


**Supplementary Figure S11. Hydrogen/Deuterium exchange analysis corroborates conformational difference between βarr1 and 2 in the lariat loop region.** Hydrogen-deuterium exchange analysis of wild-type βarr1/2 and their pre-activated, polar core mutants (βarr1$^{R169E}$βarr2$^{R170E}$) reveal significant differences in deuterium uptake between the two isoforms suggesting a conformational difference. We specifically highlight the HDX pattern observed in the lariat loop region color coded in the structural snapshot in the top panels. The original HDX data on βarrs and their pre-activated mutants have been published earlier (Yun et al., 2015), and the deuterium uptake for the peptides in the lariat loop region are re-analyzed here. The uptake plots are represented as mean±SEM of three independent experiments and analyzed using T-test (*P<0.05).
